# Transoceanic voyages of drywood termites (Isoptera: Kalotermitidae) inferred from extant and extinct species

**DOI:** 10.1101/2021.09.24.461667

**Authors:** A. Buček, M. Wang, J. Šobotník, D Sillam-Dussès, N. Mizumoto, P. Stiblík, C. Clitheroe, T. Lu, J. J. González Plaza, A. Mohagan, J. J. Rafanomezantsoa, B. Fisher, M. S Engel, Y. Roisin, T. A. Evans, R. Scheffrahn, T. Bourguignon

## Abstract

Termites are major decomposers of organic matter in terrestrial ecosystems and the second most diverse lineage of social insects. The Kalotermitidae, the second-largest termite family, are widely distributed across tropical and subtropical ecosystems, where they typically live in small colonies confined to single wood items inhabited by individuals with no foraging abilities. How the Kalotermitidae have acquired their global distribution patterns remains unresolved. Similarly, it is unclear whether foraging is ancestral to Kalotermitidae or was secondarily acquired in a few species. These questions can be addressed in a phylogenetic framework. We inferred time-calibrated phylogenetic trees of Kalotermitidae using mitochondrial genomes and nuclear ribosomal RNA genes of ∼120 species, about 27% of kalotermitid diversity, including representatives of 22 of the 23 kalotermitid genera. We found that extant kalotermitids shared a common ancestor 81 Mya (72–91 Mya 95% HPD), indicating that a few disjunctions among early-diverging kalotermitid lineages may predate Gondwana breakup. However, most of the ∼40 disjunctions among biogeographic realms were dated at less than 50 Mya, indicating that transoceanic dispersals, and more recently human-mediated dispersals, have been the major drivers of the global distribution of Kalotermitidae. Our phylogeny also revealed that the capacity to forage is often found in early-diverging kalotermitid lineages, implying that the ancestors of Kalotermitidae were able to forage among multiple wood pieces. Our phylogenetic estimates provide a platform for a critical taxonomic revision of the family and for future comparative analyses of Kalotermitidae.

## INTRODUCTION

Termites are eusocial cockroaches comprising ∼3000 described species classified into nine extant families (Krishna et al., 2013). The Termitidae, also called higher termites, are the most diverse family, including ∼2100 described species, or about three-fourth of all termite diversity (Krishna et al., 2013). By comparison, the Kalotermitidae, the second-largest family, are substantially less species-rich, including ∼450 described living species classified into 23 genera (Engel et al., 2009a; Krishna et al., 2013). However, the modest diversity of Kalotermitidae belies their considerable ecological importance. Kalotermitidae are among the most widely distributed of termite lineages, occurring worldwide between 45°N and 45°S latitude (Emerson, 1969; Jones and Eggleton, 2011). The family is also of economic significance as one-third of invasive termite species are kalotermitids, including some of the most destructive pests, such as *Cryptotermes brevis* (Evans et al., 2013). Despite their importance, there has been no attempt to reconstruct a robust and comprehensive phylogenetic hypothesis for Kalotermitidae, representing both of the taxonomic diversity and geographic breadth of the lineage. A robust phylogeny for Kalotermitidae is needed to shed light on their complex evolutionary history.

Among extant families, Kalotermitidae are one of the oldest (Inward et al., 2007; Legendre et al., 2008; Engel et al., 2009a, 2016), with fossil records dating back to the early Late Cretaceous, 99 million years ago (Mya) (Engel et al., 2007b; Barden and Engel, 2021). Time-calibrated phylogenies based on transcriptomes and mitochondrial genomes have estimated that the kalotermitid lineage diverged from their common ancestor with Neoisoptera approximately 127 Mya [96–148 Mya, combined 95% highest posterior density (HPD) of multiple analyses] (Bourguignon et al., 2015; Buček et al., 2019). These studies also estimated the age of extant (*i.e*., crown) Kalotermitidae at 69–81 Mya (54–90 Mya, combined 95% HPD of multiple analyses), albeit with limited taxon sampling within Kalotermitidae (Bourguignon et al., 2015; Buček et al., 2019). Therefore, the age of early-diverging Kalotermitidae lineages falls within the time range estimations of the final stage of Gondwanan breakup, prior to disappearance of land bridges connecting nascent continents (Upchurch, 2008). Although vicariance through continental drift may explain the distribution patterns of some early-branching kalotermitid lineages, many genera distributed across almost all terrestrial biogeographic realms, such as *Neotermes*, *Glyptotermes*, and *Cryptotermes* (Krishna et al., 2013) are far less than 60 million years old (Buček et al., 2019), postdating the breakup of Gondwana (Scotese, 2004). Therefore, both vicariance and dispersal processes have likely contributed to the geographical distribution and evolution of extant Kalotermitidae.

Termites have winged alates, yet they are poor flyers, unable to actively perform long-range dispersals (Nutting, 1969). However, the Kalotermitidae are particularly well-suited to an alternative passive dispersal strategy: transoceanic rafting. The ability of Kalotermitidae to disperse over-water is well illustrated by studies of the fauna of the Krakatau islands, which were entirely defaunated by a volcanic eruption in 1883 and then recolonized by multiple species of Kalotermitidae within 100 years (Abe, 1984; Gathorne-Hardy and Jones, 2000). The dispersal ability of Kalotermitidae stems from their lifestyle, as they usually nest in and feed on single pieces of wood (Abe, 1987; Scheffrahn et al., 2006; Scheffrahn and Postle, 2013), which can float across oceans as rafts (Thiel and Haye, 2006). The Kalotermitidae are also able to produce secondary reproductives (Myles, 1999), increasing the chance of survival of small colony fragments rafting across oceans in wood pieces, and then reproduce upon arrival to their new destination (Evans et al., 2013). These traits also predispose some species of Kalotermitidae to become invasive, spreading with the help of anthropogenic global material transport (Evans et al., 2013).

Most extant species of Kalotermitidae are unable to forage outside their nesting wood piece. Instead, they make small colonies in wood items like dead branches on living trees (Abe, 1987). In these colonies, all members retain their reproductive potential and the ability to develop wings to disperse and found new colonies, except for the soldiers that remain permanently sterile and wingless (Roisin and Korb, 2011). These life traits were intuitively interpreted as representative of the ancestral condition of modern termites (Abe, 1987; Thorne, 1997; Inward et al., 2007). By contrast, some studies have hypothesized that a complex colony organization, with true workers foraging outside their nests, is ancestral to extant termites, in which case foraging ability was secondarily lost in certain termite lineages (Morgan, 1959; Watson and Sewell, 1985; Bordereau and Pasteels, 2011; Bourguignon et al., 2016a; Mizumoto and Bourguignon, 2020). Whether foraging is ancestral to Kalotermitidae or was secondarily acquired remains unresolved.

The most comprehensive effort to reconstruct the evolutionary history of Kalotermitidae dates to the 1960s, with the taxonomic revision of Kalotermitidae by Krishna (1961). Krishna’s generic classification was based on soldier and imago morphology and has been remarkably stable since its inception, with only two new genera added (Ghesini et al., 2014; Scheffrahn et al., 2018). However, Krishna’s generic classification of Kalotermitidae has not been adequately tested by modern phylogenetic methods. A handful of studies have investigated kalotermitid phylogeny using a few genetic markers from limited taxonomic and geographic samples (e.g., Thompson et al., 2000; Ghesini et al., 2014; Bourguignon et al., 2015; Scheffrahn et al., 2018; Buček et al., 2019). Here, we used mitochondrial genomes and nuclear ribosomal RNA gene data of 173 kalotermitid samples collected worldwide, which includes species of 22 kalotermitid genera to reconstruct a robust estimate of relationships. Using this comprehensive phylogeny, we reconstructed the global historical biogeography of Kalotermitidae, identifying past natural and human-mediated dispersal events, and distribution patterns that bear the signature of potential events of vicariance through continental drift. Finally, we created a list of species of Kalotermitidae with available foraging data and discussed the evolution of such abilities according to patterns implied by the phylogenetic framework.

## RESULTS AND DISCUSSION

### Evolution of Kalotermitidae inferred from taxonomically representative phylogenetic trees

We gathered a dataset comprising the mitochondrial genomes and nuclear 18S and 28S rRNA genes of 173 samples of Kalotermitidae collected across all biogeographic realms (Figure 1). We reconstructed four phylogenies, two maximum likelihood topologies and two time-calibrated trees, inferred with and without third codon sites of protein-coding genes. For simplicity, the four trees were summarized into a reduced summary evidence tree (RSE-tree), on which the statistics of the four trees were summed (Figure 1). The RSE-tree was obtained by pruning 36 tips that diverged less than 1 Mya and were represented by samples collected in the same country. Our RSE-tree was composed of 138 tips representing 22 of the 23 genera of Kalotermitidae and including more than 120 species, which is equivalent to ∼27% of described kalotermitid species-level diversity (Krishna et al., 2013) (Figure 1). Our RSE-tree was composed of 137 branches, 119 of which were recovered in all four analyses, while 18 nodes had at least one topological conflict among analyses. The position of genus-level nodes was congruent among all four phylogenetic analyses except for 1) *Longicaputerme*s and *Kalotermes* + *Ceratokalotermes*, which were placed as the first and second diverging branches within Kalotermitidae varying in order among analyses, and 2) *Neotermes* group B and *Incisitermes* group C whose positions were variable among analyses (Figure 1).

**Figure 1.**
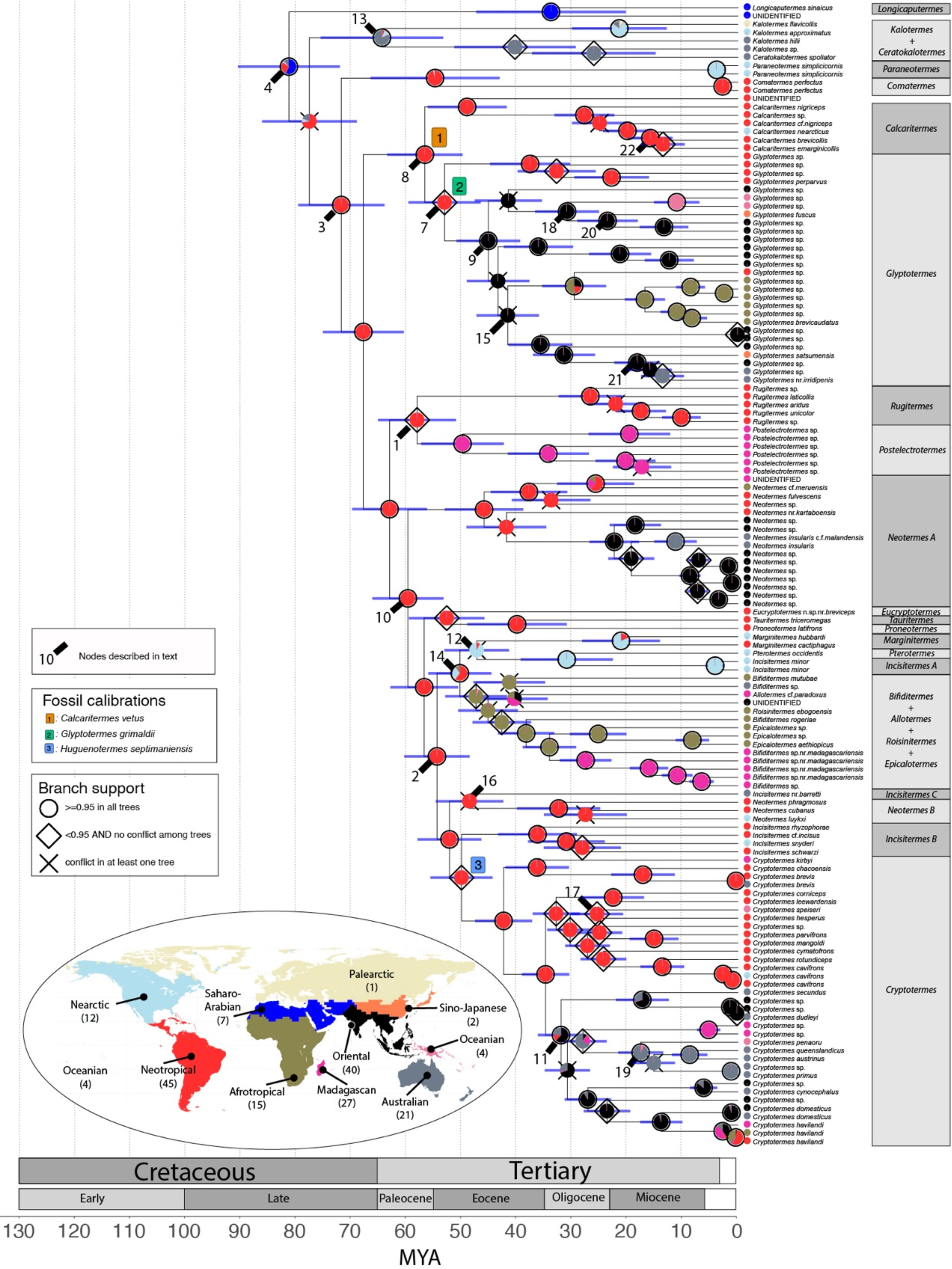
Time-calibrated phylogeny of Kalotermitidae. The tree was inferred with BEAST2 from sequence alignments excluding third codon positions. The node shapes (circle, diamond, and cross) summarize the congruence among all phylogenetic analyses. Non-kalotermitid outgroups were trimmed from the tree. Node bars represent 95% HPD intervals. Node pie charts represent the average probabilities of ancestral distributions inferred with four Bayesian Binary MCMC models. The world map indicates the biogeographic realms recognized in this study and the number of kalotermitid samples analyzed for every realm. For the phylogeny including all samples in this study and their collection codes, see Figure S1.

Time-calibrated phylogenetic trees were inferred using termite fossils as node calibrations. The timeline of termite evolution inferred with both datasets was largely congruent, with a mean node age difference of ∼5 million years (Figure S2–3). The analysis of the dataset with third codon positions included yielded older node age estimates, reflecting the high level of saturation found in third codon positions, which is known to inflate time estimates (Zheng et al., 2011). Therefore, we used the time estimates of the analysis of the dataset without third codon positions as RSE-tree (Figure 1). Our Bayesian Binary MCMC ancestral range reconstructions run on our four phylogenetic estimates with four biogeographic models yielded congruent ancestral range distributions (see Table S2–S5). These results were further corroborated by ancestral range reconstructions performed with the dispersal-extinction-cladogenesis model (Figure S5).

### Taxonomic note

Our phylogenetic trees recovered many of the currently recognized genera of Kalotermitidae as non-monophyletic. These non-natural divisions led to misinformed hypotheses on dispersal routes and ancestral distributions of Kalotermitidae (e.g., Eggleton and Davies, 2003). Therefore, our phylogenetic estimates highlight that the Kalotermitidae are in need of generic revision. Although a comprehensive revision of Kalotermitidae is outside the scope of the present paper, we provide a provisional list of genus-level taxonomic revisions that are required and carried out a handful of taxonomic changes following the International Code of Zoological Nomenclature.

*Bifiditermes* was retrieved as polyphyletic, forming a clade within which *Allotermes*, *Roisinitermes*, and *Epikalotermes* were nested. This clade needs a substantial taxonomic revision, which is outside the scope of this paper.

*Procryptotermes* was found to form two distinct lineages nested within *Cryptotermes*, a result previously found by Thompson et al. (2000) based on a mitochondrial COII phylogeny of Australian Kalotermitidae. *Procryptotermes* differs from *Cryptotermes* by the morphology of its soldiers — possessing slight differences in serration of the anterior margin of the pronotum, longer mandibles, and a less phragmotic head — while the alate imagoes of both genera are similar in all respects (Krishna, 1961). Given their considerable morphological similarities, the variability of the diagnostic characters, the existence of species with intermediary morphology (Scheffrahn and Krecek, 1999), and that both genera are mutually paraphyletic, we consider *Procryptotermes* Holmgren 1910 as a new junior subjective synonym of *Cryptotermes* Banks 1906 (new synonymy). Hereafter, we will use the generic name *Cryptotermes* to refer to both *Cryptotermes* and its junior synonym, *Procryptotermes*. Species formerly included in *Procryptotermes* are here considered formal new combinations in *Cryptotermes*: *Cryptotermes australiensis* (Gay), *C. corniceps* (Snyder), *C. dhari* (Roonwal and Chhotani), *C. dioscurae* (Harris), *C. edwardsi* (Scheffrahn), *C. falcifer* (Krishna), *C. fryeri* (Holmgren), *C. hesperus* (Scheffrahn and Křeček), *C. hunsurensis* (Thakur), *C. krishnai* (Emerson), *C. leewardensis* (Scheffrahn and Křeček), *C. rapae* (Light and Zimmerman), *C. speiseri* (Holmgren and Holmgren), *C. valeriae* (Bose).

The monospecific genus *Ceratokalotermes* was represented in our phylogeny by a single COII sequence previously published by Thompson et al. (2000). The genus was found to be nested within a paraphyletic *Kalotermes*, as originally suggested based on morphology (Hill, 1942). These results are in contrast to a previous phylogenetic reconstruction based on COII sequences in which *Ceratokalotermes* was retrieved as sister to *Kalotermes*, albeit with low support and without the inclusion of species of *Kalotermes* from outside the Australian region (Thompson et al., 2000). Although our phylogenetic estimates strongly support the paraphyly of *Kalotermes* with respect to *Ceratokalotermes*, we did not sequence any specimen of *Ceratokalotermes* and consider it premature to synonymize *Ceratokalotermes* with *Kalotermes*. The sequencing of additional samples of *Ceratokalotermes* is needed to establish its synonymy with *Kalotermes*.

The genus *Postelectrotermes* is defined by the presence of a spine on the outer margin of the mesotibia, near the outer apical spur (Krishna, 1961). We recovered a group of Madagascan kalotermitid species that includes that includes diagnostically spined *Postelectrotermes* and species that do not have this additional spine. The name *Postelectrotermes* would be available for this broader combined group, although a revision is needed to determine what morphological traits serve to circumscribe this clade of species, currently comprising *Postelectrotermes* and unplaced Malagasy species erroneously assigned to *Glyptotermes*.

*Incisitermes* is a polyphyletic genus containing three distantly related lineages: *Incisitermes* group A includes the Nearctic *Incisitermes minor*; *Incisitermes* group B includes Nearctic and Neotropical species; and *Incisitermes* group C contains the Australian *Incisitermes* nr. *barretti*. Each of these three groups presently forming *Incisitermes* diverged from their sister group 30– 50 Mya and should be classified into three separate genera. We did not sequence the type species of the genus, *I*. *schwarzi*, but morphological characters should be sufficient to properly identify the groups and determine which clade comprises *Incisitermes* proper and which represent new genera. For example, *I. schwarzi*, *I. minor*, and *I. marginipennis* all have soldiers with pigmented compound eyes, at least suggesting that *Incisitermes* A represents that clade to which the generic name *Incisitermes* should be restricted (with *Incisitermes* B therefore likely representing a new genus). At the least, *Incisitermes* C assuredly represents an unnamed genus. It is possible that the principally Asiatic and Philippine species (e.g., *I. laterangularis*, *I*. *inamurai*, and *I*. *didwanaensis*) also represent a distinct genus (Engel, pers. obs.), although we were not able to test this in our analyses. *Incisitermes peritus* in Dominican amber likely represents clade A, while *I. krishnai* in Mexican amber could represent clade B, but these will require critical testing once these clades are formally revised.

*Neotermes* is also a polyphyletic assemblage represented in our phylogenies by two distant groups, *Neotermes* A and *Neotermes* B. Both *Neotermes* groups share their last common ancestor 60 Mya (53–66 Mya 95% HPD, node 10 in Figure 1). The two *Neotermes* groups should be classified into two separate genera. At least based on preliminary comparisons, the type species, *N*. *castaneus*, is more closely allied to those species in *Neotermes* B (Engel and Krishna, pers. obs.), implying that those South American, Oriental, and Australian species of *Neotermes* A represent an unnamed genus.

### Historical biogeography of kalotermitids

We estimated the divergence between the lineages comprising Kalotermitidae and Neoisoptera at 111 Mya (100–122 Mya 95% HPD, Figure S1). Although this divergence age estimate is younger than that estimated by previous time-calibrated phylogenetic trees using mitochondrial genomes and transcriptome data, it remained congruent, as the HPD intervals largely overlapped (Bourguignon et al., 2015; Buček et al., 2019). These differences may result from using different fossil calibrations among studies, by changes of fossil age estimates based on revised or refined stratigraphic and radiometric dating for particular fossil deposits, or by different taxonomic sampling among studies. Regardless, all studies estimated the origin of Kalotermitidae to predate the first kalotermitid fossils, the 99-million-year-old species of *Proelectrotermes* (Engel et al., 2007b), by 6 to 28 million years. Crown-group Kalotermitidae share a common ancestor estimated to live 81 Mya (72–91 Mya 95% HPD, node 4 in Figure 1).

Given that the common ancestor of modern Kalotermitidae is estimated at 81 Mya (72–91 Mya 95% HPD), vicariance may explain the distribution of early-diverging kalotermitid lineages. Our analyses did not allow us to identify the ancestral range of the last common ancestor of kalotermitids. However, four biogeographic realms, the Saharo-Arabian, Neotropical, Australian, and Nearctic, were inferred as possible ancestral ranges (Figure 1). There is also a possibility that the ancestor of modern Kalotermitidae was distributed across several biogeographic realms, such as the Neotropical and Australian realms that were connected via land bridges, such as the Weddellian Isthmus, until at least ∼56 Mya (Reguero et al., 2014). While the ancestral range of the last common ancestor of modern Kalotermitidae remains speculative, we found strong support for a Neotropical origin of the clade composed of all modern Kalotermitidae with the exclusion of the three species-poor genera *Kalotermes*, *Ceratokalotermes*, and *Longicaputermes*. The age of this clade was estimated at 72 Mya (64–80 Mya 95% HPD, node 3 in Figure 1), which predates the separation of Antarctica from South America and Australia (Reguero et al., 2014). Admittedly, at the time of the estimated age of this clade, the distinction of a Neotropical region as unique from the remainder of the Gondwanan landmasses was less meaningful owing to the lack of full paleogeographical differentiation at that time and it remains unknown to what degree extinction has impacted crown-Kalotermitidae. Nonetheless, it is safe to at least conclude that among modern species those in the Neotropical region appear to be more relict than those of other realms, which by contrast are nested in more derived clades (Figure 1). Notably, one of the two potential sister groups of this clade, the group *Kalotermes* + *Ceratokalotermes*, possibly has an Australian origin, in which case vicariance through continental drift provides an explanation for the ancestral distribution of these two early-diverging kalotermitid lineages. However, this scenario remains speculative and requires further testing. Other disjunctions between biogeographic realms postdated the breakup of Gondwana and are therefore most likely explained by transoceanic and land dispersals rather than by vicariance (Figure 1).

Our results indicate that the Neotropical realm is likely a cradle of modern kalotermitid diversity and the source of repeated dispersals to other biogeographic realms (Figures 1–2). These results were not a consequence of our more intensive sampling of the Neotropical realm because our ancestral range reconstructions were robust to the removal of up to 2/3 of the Neotropical samples (Figures S6–7). We also found evidence for two natural long-distance over-water dispersal events to the Neotropical realm: 1) *Glyptotermes* dispersed to the Neotropics from the Afrotropical or Oriental realm during the Eocene, 42 Mya (36–47 Mya 95% HPD, node 15 in Figure 1), and 2) *Marginitermes* dispersed to the Neotropics from the Nearctic realm 47 Mya (42–53 Mya 95% HPD, node 12 in Figure 1). Note that throughout this work we date dispersals using the maximum dispersal age, that is, the age of the youngest hypothesized ancestor, which has a different realm than its extant descendants (see Figure 2 and Methods for additional details) with >90% probability. Other kalotermitid dispersal events to the Neotropics were mediated by human activities with notable introductions of *Cryptotermes havilandi* from the Afrotropical and *C. dudleyi* from the Oriental realm (Evans et al., 2013). One possible explanation for this observed low dispersal rate to the Neotropics is the presence of a diverse neotropical kalotermitid fauna, which presumably filled most niches available to Kalotermitidae, hindering colonization by foreign kalotermitids.

**Figure 2.**
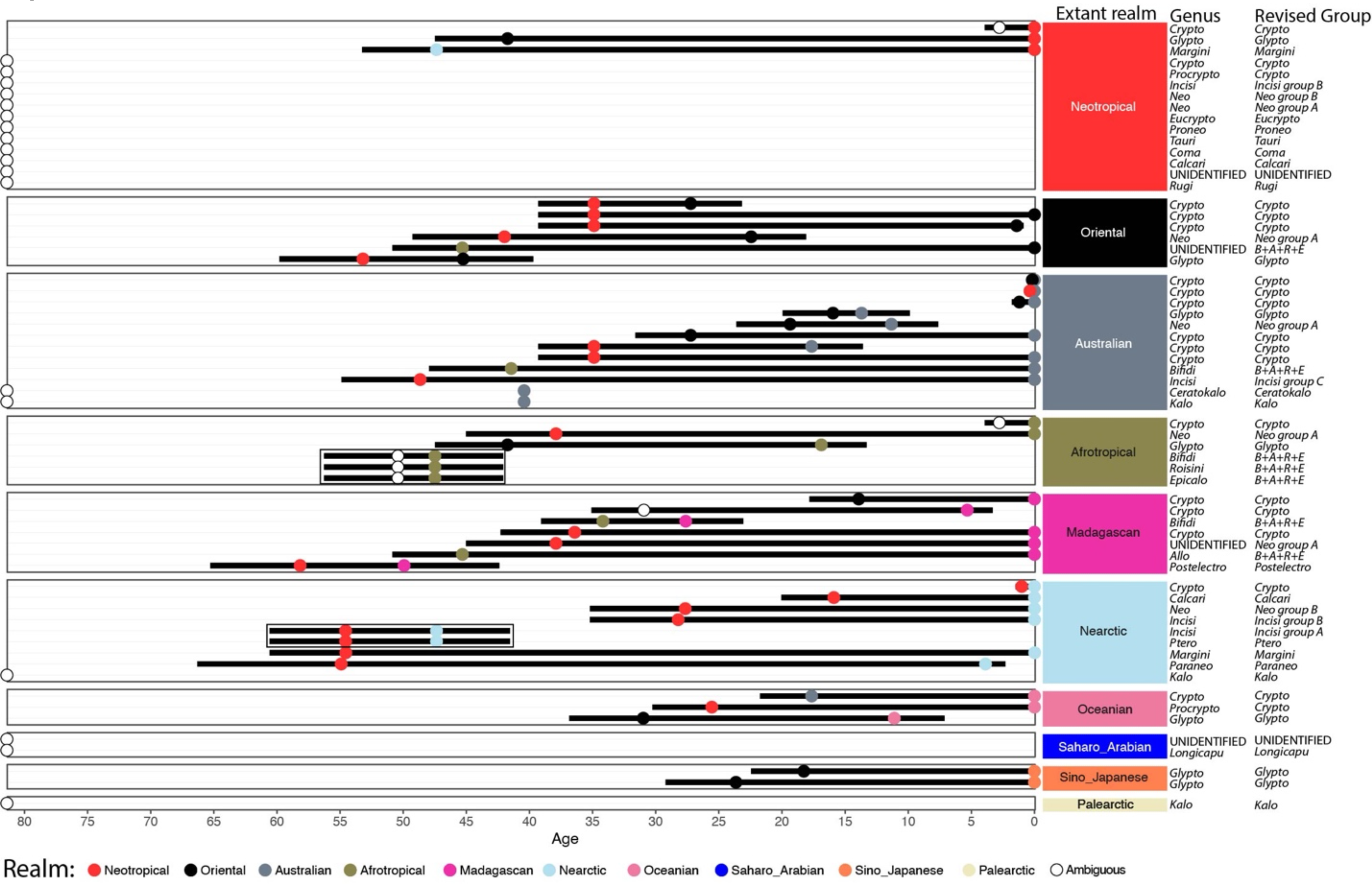
Dates of Kalotermitidae dispersal to their extant realms. The black bars indicate the age intervals as combined 95% HPD intervals of 1) the youngest ancestor of extant genera that with high probability (>90%) did not share realm with them and 2) the oldest representative of extant genera that with high probability (>90%) diversified sympatrically within the realm. The dispersal age intervals are plotted separately for each genus and each realm. For genera represented by a single species the oldest representative is the extant species. All ancestor nodes with single realm probability >90% are colored according to their high-probability realm. Ancestor nodes with ambiguous realm (<90% realm probability) are indicated by empty circle. Empty circle positioned arbitrarily at the age of the crown-Kalotermitidae indicate the genera for which no high probability realm disjunctions were inferred. The groups of genera that diversified sympatrically within the extant realm and thus represent a single dispersal event are framed in rectangle. The generic names were abbreviated for clarity by excluding the “*termes*” suffix when possible. “B+A+R+E” indicates a monophyletic group encompassing *Bifiditermes*, *Allotermes*, *Roisinitermes*, and *Epicalotermes*. All dates are extracted from Figure 1.

The Oriental realm was colonized by at least four dispersal events (Figures 1–2). Our ancestral range reconstructions indicate that *Glyptotermes* was the first lineage dispersing to the Oriental realm in the Early Eocene, 53 Mya (47–60 Mya 95% HPD, node 7 in Figure 1). Subsequent dispersal events to the Oriental realm took place 35–45 Mya (31–51 Mya combined 95% HPD), and prior to the Eocene-Oligocene climatic shift and extinction events. These dispersal events included one dispersal of *Neotermes* group A and one or several dispersals of *Cryptotermes*, all of which originated from the Neotropical realm (Figures 1–2). The Oriental realm was also colonized by one additional lineage, wrongly identified as *Neotermes* by Bourguignon et al. (2015), and allied to the *Bifiditermes* + *Allotermes + Roisinitermes* + *Epicalotermes* group, which most likely dispersed from the Afrotropics (Figure 1). Although our time-calibrated trees suggest the first Oriental kalotermitids date from the early Eocene, the fossils of *Proelectrotermes* provide evidence that the Oriental realm was already occupied by kalotermitids >99 Mya (Engel et al., 2007b). Note that *Proelectrotermes* could belong to stem-kalotermitids and outside of the clade of crown-Kalotermitidae. Our ancestral range reconstructions do not support an Oriental origin for crown-kalotermitids, and analyses with the last common kalotermitid ancestor constrained to be Oriental supported that the modern Oriental kalotermitid fauna colonized the Oriental realm through at least four dispersals, including at least three long-distance over-water dispersals (Figure S5). Early kalotermitid fossils unearthed from the Oriental realm, such as the 99-million-year-old Kachin amber fossils of *Proelectrotermes*, therefore belong to ancient lineages that went extinct in the Oriental realm. If *Proelectrotermes*, perhaps along with concurrent species of *Kachinitermes* and *Kachinitermopsis*, is representative of stem-kalotermitids, then the broader lineage encompassing Kalotermitidae could have originated outside of the Neotropical realm, with crown-kalotermitids representing an originally “western Gondwana” subordinate clade of this broader group. Refined testing of such a hypothesis will require significantly more Cretaceous fossils of stem-Kalotermitidae and a phylogeny ascertaining their relationships to one another as well as to the crown group. At the least, there is no current phylogenetic or paleontological evidence to support an eastern Laurasian (i.e., north of Tethys) origin for the original divergence from the common ancestor of Neoisoptera.

Madagascar was colonized by kalotermitids at least seven times independently (Figures 1–2). The first dispersal event to Madagascar was that of the ancestor of the Madagascan *Postelectrotermes*, which diverged from the common ancestor of Neotropical *Rugitermes* 58 Mya (51–65 Mya 95% HPD, node 1 in Figure 1). Two more colonization events were of Neotropical origin, that of *Neotermes* group A 38 Mya (31–45 Mya 95% HPD) and that of the lineage comprising *Cryptotermes kirbyi* 36 Mya (31–42 Mya 95% HPD), and one colonization event from the Afrotropical realm, that of *Bifiditermes* 34 Mya (29–39 Mya 95% HPD). The origins of the three other dispersal events to Madagascar are uncertain. *Cryptotermes havilandi* is presumably invasive in Madagascar (Evans et al., 2013). But divergence of Madagascan *C. havilandi* and non-Madagascan *C. havilandi* dating to 3 Mya (2–4 Mya 95% HPD) suggests its dispersal to Madagascar might predate human voyages. More thorough sampling in Afrotropical realm will be necessary to resolve the native range of *C. havilandi*. *Allotermes* colonized Madagascar from the Afrotropical or Oriental realms 45 Mya (40–51 Mya 95% HPD) and the third lineage of *Cryptotermes* colonized Madagascar 31 Mya (27–35 Mya 95% HPD) from either the Oriental or Australian realm (Figure 1). Madagascar has been isolated from other landmasses since at least ∼140 Mya, first connected to India after both regions broke away from Africa, then alone after India broke apart from Madagascar and started drifting northward ∼85 Mya (Gibbons et al., 2013). The long isolation of Madagascar from other biogeographic realms implies that Kalotermitidae colonized Madagascar repeatedly through long-distance over-water dispersals.

Australia was colonized multiple times by Kalotermitidae. Our ancestral range reconstructions suggest the last common ancestor of Kalotermitidae could possibly be of Australian origin, and, if so, then it is unclear whether the clade composed of *Kalotermes* and *Ceratokalotermes* dispersed to Australia, or whether their presence in Australia is relict. The timing of other disjunctions between the Australian and other realms correspond to natural dispersal events or human introductions. Our ancestral state reconstructions retrieved four colonization events of Australia by non-*Cryptotermes* Kalotermitidae and up to six colonization events by *Cryptotermes* (Figure 1). *Glyptotermes* and *Neotermes* group A dispersed over-water to Australia from the Oriental realm 16 Mya (12–20 Mya 95% HPD) and 19 Mya (15–24 Mya 95% HPD), respectively. This timing coincides with the colonization of Australia by higher termites, which possibly benefited from the Miocene expansion of grasslands and growing drier climates in Australia (Bourguignon et al., 2017). The other two colonization events by non-*Cryptotermes* Kalotermitidae were more ancient. *Bifiditermes*, which was incorrectly identified by Cameron et al. (2012) as *Neotermes insularis*, colonized Australia from Africa through long-distance over-water dispersal 41 Mya (35–48 Mya 95% HPD) and the Australian *Incisitermes* group C, which includes *Incisitermes* nr. *barretti*, colonized Australia from the Neotropics by land or through over-water dispersal 49 Mya (43–55 Mya 95% HPD, node 16 in Figure 1). Because we did not include the Oriental “*Incistermes*” in our sampling, it remains to be determined whether its inclusion would alter this reconstruction. Thus, the historical biogeographic origin of Australian “*Incisitermes*” requires further testing. South America and Australia were connected by land through Antarctica until at least ∼56 Mya (Reguero et al., 2014). The formation of deep-water isolation by the Eocene-Oligocene transition, when the Antarctic circumpolar current was established, resulted in an eventual permanent glaciation of Antarctica and a global cooling and drying during the latest Paleogene and early Neogene (Zachos, 2001). *Incisitermes* group C therefore dispersed either over-water or by land, as was previously suggested for many animal taxa, including insects (Sanmartín and Ronquist, 2004), which possibly dispersed from South America to Australia through an Antarctic landmass covered by temperate forests (Dawson et al., 1976; Mörs et al., 2020). In addition, *Marginitermes absitus*, which was absent from our phylogenetic inferences, presumably colonized northern Australia via a long-distance over-water dispersal event from Central or North America (Scheffrahn and Postle, 2013). Our ancestral range reconstructions indicate up to six colonization events of Australia by *Cryptotermes*, although this number was uncertain because of ambiguous ancestral range reconstructions (node 11 in Figure 1). Four of these six colonization events correspond to human introductions, from the Oriental realm in the case of *C. domesticus*, *C. dudleyi*, and *C. cynocephalus*, and from the Neotropical realm in the case of *C. brevis* (Evans et al., 2013). The origin of the two other colonization events, that of *C. secundus* and the clade of *Cryptotermes* including *C. austrinus*, is likely the Oriental realm but we cannot definitively exclude a Neotropical origin. Future studies are required to refine these patterns of dispersals, especially for *Cryptotermes*.

Our ancestral range reconstructions identified four independent colonization events of the Afrotropical realm by Kalotermitidae (Figures 1–2). The most ancient dispersal event was that of *Bifiditermes* + *Epicalotermes* + *Roisinitermes* + *Allotermes*, which colonized the Afrotropical realm through long-distance over-water dispersal from the Neotropical or Nearctic realms 50 Mya (45–56 Mya 95% HPD, node 14 in Figure 1). Two more long-distance over-water dispersals to the Afrotropical realm involved *Glyptotermes* and *Neotermes*, the former originated from the Oriental or Neotropical realm and dispersed 42 Mya (36–47 Mya 95% HPD, node 15 in Figure 1), while the latter originated from the Neotropical or Madagascan realm and dispersed 38 Mya (31–45 Mya 95% HPD). Lastly, *Cryptotermes havilandi* is an invasive species believed to be native to West Africa (Evans et al., 2013), in which case its lineage colonized the Afrotropical realm from the Oriental realm 14 Mya (10–18 Mya 95% HPD). Alternatively, *C. havilandi* was introduced to the Afrotropical realm by human activities from an unknown location (Figure 1). Therefore, the Afrotropics might have been naturally colonized by Kalotermitidae only thrice independently. This could possibly reflect the abundance of higher termites, whose center of origin is the Afrotropical realm (Bourguignon et al., 2017), and which perhaps hindered the colonization of the realm by kalotermitids.

Our sampling of the Oceanian, Sino-Japanese, Saharo-Arabian, and Palearctic realms was limited, although it somewhat reflects the low termite diversity in these realms. Most notably, the Saharo-Arabian realm is inhabited by one of the earliest-branching kalotermitid lineages, represented in this study by *Longicaputermes* and an unidentified species from which we only collected immature individuals. Therefore, despite its low termite diversity, the Saharo-Arabian realm is either a potential center of kalotermitid origin, or was invaded by early diverging kalotermitids. Naturally, modern distributions do not necessarily reflect the ancient distribution of a clade as evidenced by many fossils that demonstrate the historical ancestral distribution of a clade was elsewhere from surviving species of that same clade (e.g., honey bees: Engel et al., 2009; Kotthoff et al., 2013; megalyrid wasps: Vilhelmsen et al., 2010; thorny lacewings: Nakamine et al., 2020; mastotermitid termites: Krishna et al., 2013; and many other examples in Grimaldi and Engel, 2005). Thus, additional sampling from sparsely surveyed Saharo-Arabian realm is necessary to resolve the early historical biogeography of kalotermitids since it clearly includes relict species.

Our ancestral state reconstructions indicate that the Oceanian realm was colonized at least thrice: once from the Neotropics by a lineage survived by modern *C. speiseri* 26 Mya (21–30 Mya 95% HPD, node 17 in Figure 1), once from the Oriental realm by *Glyptotermes* 31 Mya (25– 27 Mya 95% HPD, node 18 in Figure 1), and once by a lineage survived by *C. penaoru* 18 Mya (14–22 Mya 95% HPD, node 19 in Figure 1) from the Australian realm (Figures 1–2). The Sino-Japanese realm was colonized at least twice independently from the Oriental region by lineages including *Glyptotermes fuscus* 24 Mya (18–29 Mya 95% HPD, node 20 in Figure 1) and *G. satsumensis* 18 Mya (14–22 Mya 95% HPD, node 21 in Figure 1). Additional sampling from these poorly sampled realms is required to draw a more accurate picture of these aspects of kalotermitid historical biogeography.

The Palearctic realm only hosts 18 described species (Krishna et al., 2013), but its rich fossil record indicates kalotermitid presence by at least the Early Eocene (Figure S4). The absence of early-branching Palearctic kalotermitid lineages other than *Kalotermes* in our sampling (Figures 1–2) either reflects an undersampling of extant Palearctic species or, more likely, the extinction of lineages once abundant during the past global climatic optimum.

We inferred 6–7 dispersal events to the Nearctic realm, six from the Neotropics (Figures 1–2), which are consistent with the geographic vicinity of both realms since at least the Paleocene, ∼66–56 Mya (Buchs et al., 2010). The most recent colonization event of the Nearctic realm by a Neotropical kalotermitid was that of *C. cavifrons* 1 Mya (0.6–1.5 Mya 95% HPD). The five other dispersal events, involving lineages including *I. snyderi*, *Neotermes luykxi*, *Pterotermes* + *Incisitermes* group A, *Calcaritermes nearticus*, and *Paraneotermes simplicornis*, took place between 16 Mya (12–20 Mya 95% HPD, node 22 in Figure 1) and 55 Mya (49–61 Mya 95% HPD, node 2 in Figure 1), and therefore involved over-water dispersals. *Marginitermes* either diversified within the Neotropical realm from a common ancestor with *Pterotermes* and *Incisitermes* group A, or it dispersed from the Neotropical realm, thus representing a possible seventh dispersal to the Nearctic realm (Figure 1). The origin of Nearctic *Kalotermes* is more challenging to trace. *Kalotermes* either dispersed to the Nearctic realm from an unidentified location, the lineage has experienced considerable extinction, or the Nearctic realm was the center of kalotermitid origin. If the last hypothesis were the case, then the presence of the lineage that gave rise to *Kalotermes* (the age of crown-*Kalotermes* is uncertain) in the Nearctic realm dates back to earliest history of Kalotermitidae. Note that this scenario received a low probability by our ancestral range reconstructions (Figure 1). The modern kalotermitid fauna of the Nearctic realm is overwhelmingly of Neotropical origin.

### Reconciliation of extant and extinct kalotermitid evidence

Molecular phylogenies are reconstructed using extant taxa. The exclusion of extinct species can lead to spurious historical biogeographic reconstructions (Lieberman, 2002). Although fossil evidence has been integrated into biogeographic reconstructions inferred from molecular phylogenies by previous studies (e.g. Meseguer et al., 2015), these methods are hampered for a family with a fragmentary fossil record and containing many species of uncertain taxonomic affiliations like Kalotermitidae. Kalotermitids are represented by 74 fossils belonging to twelve extinct and six extant genera in the paleontological database PaleoDB (Figure S4). Despite the paucity of available fossils, the age and biogeographic origin of kalotermitid fossils are somewhat consistent with our phylogenetic inferences (nodes 7, 12, 13, 14, and 15 in Figure 1), although the earliest kalotermitids are from the Oriental rather than the Neotropical region. Indeed, the available fossil record suggests a more intricate historical biogeography for Kalotermitidae, characterized by considerable past extinctions, consistent with the fact that most organisms that have ever lived are today extinct and so historical biogeographic patterns must always be tempered by a consideration of significant extinction. Extinction events are at minimum evidenced by at least four groups of Kalotermitidae: 1) *Electrotermes* is the first unambiguous kalotermitid fossil from the Palearctic realm represented by ∼49–41 million-year-old fossils (Nel and Bourguet, 2006), and predating our oldest estimates of Palearctic kalotermitids (Figure 1 and S4); 2) *Huguenotermes*, an extinct genus allied to modern *Cryptotermes*, which appeared in the Palearctic at least by 35 Mya (Engel and Nel, 2015), while there are no modern native species of *Cryptotermes* or related genera in the Palearctic realm; 3) a wholly unique fauna of kalotermitids from the Miocene of Zealandia which may have diverged from other Kalotermitidae when Zealandia separated from Australia in the Late Cretaceous, and then become extinct as the landmasses of Zealandia became submerged during the Neogene, necessitating more recent recolonization of modern New Zealand (Engel and Kaulfuss, 2017); and 4) the oldest kalotermitid fossils of *Proelectrotermes* (a genus also represented in 41 Ma Baltic amber of the Palearctic realm), *Kachinitermes*, and *Kachinitermopsis* from the ∼99 million year old Kachin amber (Engel et al., 2007b), substantially predating our estimates of the oldest Oriental kalotermitids (53 Mya, 47–60 Mya 95% HPD, node 7 in Figure 1). These last three are possible representative of the kalotermitid stem group, but the others are crown-Kalotermitidae. These fossils imply the extinction of ancient kalotermitid faunas, or subsets of those faunas, in the Palearctic and Oriental realms, which, in the case of at least *Proelectrotermes*, *Kachinitermes*, and *Kachinitermopsis*, involved the extinction of stem-kalotermitids (Engel et al., 2009a, 2016). Furthermore, extinct representatives of many other kalotermitid genera (Krishna et al. (2013), and Figure S4) provide the smallest of glimpses of localized extinctions throughout the phylogeny of Kalotermitidae.

### Dispersal potential and invasive traits

Herein, we explored kalotermitid phylogeny based on the most comprehensive taxonomic sampling yet undertaken, including most major lineages sampled across their distribution. We did not attempt to include samples of invasive species from across their introduced range and, therefore, many recent human-assisted kalotermitid dispersals are not represented in our ancestral range reconstructions (Evans et al., 2013). Altogether, we inferred ∼40 kalotermitid dispersals among biogeographic realms, most of which were long-distance oceanic dispersals. In comparison, our previous study on the historical biogeography of Termitidae based on a representative taxonomic sampling of 415 species (∼18% of termitid species diversity) revealed about 30 dispersals among realms (Bourguignon et al., 2017). Our previous phylogeny of the clade *Heterotermes* + *Coptotermes* + *Reticulitermes*, based on 37 of the 258 extant species, inferred 14 dispersals among biogeographic realms (Bourguignon et al., 2016b), and our phylogeny of Rhinotermitinae, based on 27 of 66 described species, retrieved four dispersals among biogeographic realms (Wang et al., 2019). The high number of long-distance over-water dispersals inferred for kalotermitids is consistent with their lifestyle, suitable for transoceanic voyages in rafting wood.

Not all branches of the kalotermitid tree had comparable dispersal success. One branch with high-dispersal success is that of *Cryptotermes* (Figure 1), a genus that also includes some of the most successful invasive termite species, and the most successful invasive species of Kalotermitidae (Evans et al., 2013). *Cryptotermes* includes five invasive species, *C. havilandi*, *C. cynocephalus*, *C. dudleyi*, *C. brevis*, and *C. domesticus*, making it one of the most cosmopolitan of termite genera (Evans et al., 2013). Species of *Cryptotermes* possess three biological traits that enhanced their invasiveness and have allowed them to disperse around the world with human assistance: they are wood-feeders, nesting in wood pieces, and easily produce secondary reproductives (Evans et al., 2013). In addition, *C. domesticus* is tolerant to saline water, a biological trait that presumably increases its ability to survive long-distance oceanic dispersal in wood pieces and possibly human-assisted journeys (Chiu et al., 2021). Additional life history traits might contribute to the invasiveness of *Cryptotermes*, but the physiology and ecology of Kalotermitidae have not been fully elucidated, preventing us from comprehending what may also aid their suitability for such passive dispersal. This question could be adressed using a taxonomically representative sampling of kalotermitids. The phylogenies of Kalotermitidae reconstructed here provide a framework to guide such sampling.

### Evolution of social organization

Most extant species of Kalotermitidae are unable to forage outside their nests. This life trait has been alternatively considered as either ancestral for termites (Abe, 1987; Thorne, 1997; Inward et al., 2007), or as a derived condition stemming from the secondary loss of foraging abilities in some lineages such as kalotermitids (Morgan, 1959; Watson and Sewell, 1985; Bordereau and Pasteels, 2011; Bourguignon et al., 2016a; Mizumoto and Bourguignon, 2020). Our taxonomically representative phylogeny of Kalotermitidae represents a starting point to disentangle the evolution of social organization in Kalotermitidae.

Due to the scarcity of behavioral observations, we do not attempt to reconstruct the ancestral behavior of Kalotermitidae quantitatively. However, some pieces of information on the behavior of Kalotermitidae are available for several species. Some Kalotermitidae are known to have the ability to build underground tunnels and shelter tubes covering foraging trails. This is notably the case of the two early branching kalotermitid lineages (Figure 3 and Table S8). Indeed, *Longicaputermes* can build shelter tubes, and *Paraneotermes* builds underground tunnels to colonize new wood resources (Light, 1937) (Figure 3 and Table S8). These foraging abilities are not unique to early branching Kalotermitidae and have also been observed in other kalotermitids. For example, we observed *C. brevis* building shelter tubes under laboratory conditions (Figure 3), and *Neotermes* have been reported to build below-ground tunnels (Thompson, 1934; Goetsch, 1936; Waterhouse, 1993). Moreover, species from across kalotermitid phylogeny produce and follow trail pheromones which are generally presumed to serve termites as orientation clues outside of nest (Sillam-Dussès et al., 2009) (Figure 3). The occurrence of foraging behavior in early-branching kalotermitid lineages and in species spread across the kalotermitid phylogeny at least initially suggests that the last common ancestor of Kalotermitidae was capable of foraging among multiple wood pieces.

**Figure 3.**
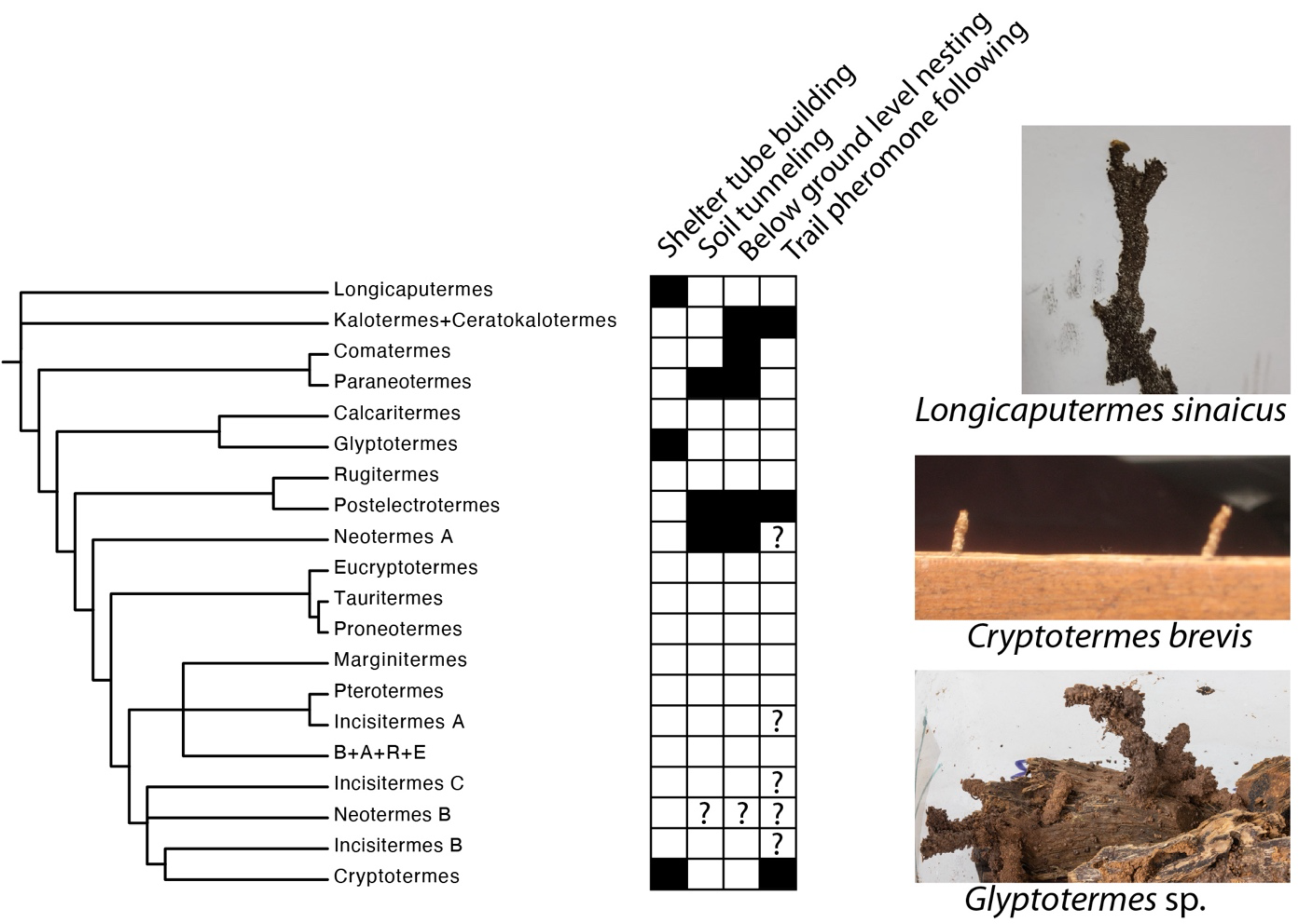
Construction and foraging abilities of Kalotermitidae. The phylogeny from Figure 1 was collapsed at the genus level. Branches showing conflicting topology among our phylogenetic analyses are represented as polytomies. The heatmap indicates the presence of behavioral record (black) for at least one species of the genus. Note that the absence of evidence of a behavior (white) either reflects its actual absence or the absence of its observation. “Below ground level” indicates presence of termites in wood items that are below ground level, such as tree roots. Interrogation marks indicate observations that cannot be unambiguously mapped onto the phylogeny due to the uncertain phylogenetic position of observed species. The photos show the shelter tubes of *Longicaputermes sinaicus* attacking timber in urban environment, the shelter tubes built by *Cryptotermes brevis* under laboratory conditions, and the shelter tubes built by an unidentified *Glyptotermes* species maintained in a laboratory and collected in Papua New Guinea. For details on literature survey of kalotermitid behavior, see Table S8.

Although several species of Kalotermitidae have foraging abilities, especially those belonging to early branching lineages, most modern species of Kalotermitidae appear to lack foraging abilities or exhibit such behaviors only under specific circumstances such as in laboratory colonies. Such facultative foraging would be consistent with an ancestral presence of the ability that is then not utilized in most lineages under natural environments, perhaps only expressing such behaviors under extreme conditions, much like facultative sociality in halictine bees (Eickwort et al., 1996). We hypothesize that the shift in kalotermitid social organization and foraging abilities was driven by competition. In particular, two lineages of Neoisoptera, *Heterotermes* + *Coptotermes* + *Reticulitermes* and the higher termites (Termitidae), are major competitors of kalotermitids owing to the overlap of their niches, high abundance, mutually high aggression, and the existence of adaptive avoidance mechanisms (Thorne and Haverty, 1991; Evans et al., 2009). Biotic factors, such as species competition, can shape species spatial distribution at a global scale (Wisz et al., 2013). Kalotermitids expressing foraging abilities often inhabit biomes hosting few or no other termite species, such as arid regions (*Longicaputermes*, *Paraneotermes,* and *Neotermes chilensis*), remote islands (*Neotermes rainbowii*) (Waterhouse, 1993), and high-elevation mountains (*Postelectrotermes militaris*, *Comatermes perfectus*) (Hemachandra et al., 2014; Scheffrahn, 2014; Gnanapragasam, 2018). This geographical distribution is consistent with foraging kalotermitids being competitively excluded from regions where wood-feeding Neoisoptera are abundant, as well as the notion that expression of foraging is only under atypical environmental conditions. As noted, kalotermitids exhibit behavioral flexibility with respect to foraging behavior as demonstrated by *Cryptotermes*, which is capable of building galleries in laboratory conditions (Figure 3). Foraging in Kalotermitidae is likely a complex interplay between extreme conditions, local competition, and perhaps yet unidentified factors. It is unclear to what extent the foraging behavior of kalotermitids could decrease their fitness in biomes occupied by competing wood-feeding Neoisoptera. Ants represent another hyper-diverse and abundant organismal group that interact with termites, mostly through predation (Wilson, 1971). Ant predation thus likely represent another factor favoring confinement of termites into a single piece of wood. Additional work is needed to quantify competition among termites and reconcile kalotermitid historical biogeography with that of their presumed wood-feeding termite competitors and ant predators.

## Conclusion

We present the current most comprehensive molecular phylogenetic reconstruction of Kalotermitidae, based on a taxonomic sampling of global kalotermitid diversity and distribution. This study is also the first to provide an estimated timeline of kalotermitid evolution and specifically reconstructions of historical biogeographic events across the family. Our ancestral range reconstructions imply that the Neotropical realm is a cradle for crown-group kalotermitid diversity, with the exception of two early diverging lineages and putative stem-group fossils from the Oriental region. We show that the distribution of modern Kalotermitidae includes upwards of 40 disjunctions between biogeographic realms, most of which can be explained by long-distance transoceanic dispersal events. While the single-piece nesting lifestyle has been sometimes considered as representative of the proto-termite colony, or at least ancestral to Kalotermitidae, we found that many kalotermitids, including early diverging species-poor genera, are capable of foraging outside their nesting piece of wood. These results suggest that the social organization of the colonies of the common ancestor to all modern Kalotermitidae was more complex than previously appreciated, and perhaps includes some degree of facultative expression. We postulate that single-piece nesting is a derived trait that evolved as a specialization to minimize niche overlap with competitors, such as species of Neoisoptera. Finally, our molecular phylogeny reveals that several kalotermitid genera are non-monophyletic and provides a framework for a future taxonomic revision of the family. At a minimum we formally synonymize the genus *Procryptotermes* under *Cryptotermes* (new synonymy) and transfer the species of the former to the latter.

## METHODS

### Sample collection

The termite samples used in this study were collected during the last three decades by the authors. Two sampling procedures were employed. Samples were either collected in 80% ethanol and stored at room temperature or collected in RNA-later® and stored at -80°C until DNA extraction.

### Mitochondrial genome sequencing, assembly, and annotation

We used different approaches for the samples preserved in RNA-later® and 80% ethanol. For the 108 samples preserved in RNA-later®, we extracted total genomic DNA from one to ten termite immatures (pseudergates) heads using the DNeasy Blood & Tissue Kit (Qiagen). The total extracted DNA was used to amplify mitochondrial genomes with TaKaRa LA Taq in two long-range PCR reactions. The primers and conditions of the reactions were as described by Bourguignon et al. (2015). The concentration of the two long-range PCR amplicons was measured with Qubit 3.0 fluorometer, and both amplicons were mixed in equimolar concentration. One DNA library was then prepared for each sample separately. Libraries were either prepared in-house, using unique dual index barcodes, or by a sequencing company, using a unique combination of index barcodes. In-house libraries were prepared with the NEBNext Ultra II DNA Library Prep Kit (New England Biolabs) using one-fifteenth of the reagent volume indicated in the manufacturer protocol. Libraries were pooled and paired-end sequenced on the Illumina HiSeq platform or the Illumina MiSeq platform. For the 57 samples preserved in 80% ethanol, total genomic DNA was extracted from entire pseudergate bodies using the DNeasy Blood & Tissue Kit (Qiagen). Libraries were prepared for shotgun sequencing, directly from whole pseudergate body DNA, without any long-range PCR step. Because the DNA extracted from these samples was generally highly fragmented, we prepared the libraries using a modified protocol. Libraries were prepared with the NEBNext Ultra II DNA Library Prep Kit (New England Biolabs) without an enzymatic fragmentation step. In addition, library preparation was downsized to one-fifteenth of the volume indicated in the manufacturer protocol for all reagents. Other steps were performed as suggested by the manufacturer protocol. One library with unique dual index barcodes was prepared for each sample separately. A total of 3 μl of each library was pooled and sequenced on the Illumina HiSeq X platform.

We obtained an average of 52 megabases and 2 gigabases of raw sequencing data for samples sequenced with the amplicon-based approach and samples sequenced with the shotgun sequencing approach, respectively. Raw sequencing data was quality-checked with FastQC (https://www.bioinformatics.babraham.ac.uk/projects/fastqc/), and trimmed and quality-filtered with bbduk (sourceforge.net/projects/bbmap/) using the following parameters: “ktrim=r k=23 mink=7 hdist=1 tpe tbo maq=10 qtrim=rl trimq=15 minlength=35”. Multiple mitochondrial genome assembly approaches were used as described below, and the longest mitochondrial genome contigs or scaffolds were selected for each sample. The mitochondrial genomes obtained through long-range PCR amplifications were assembled using SPAdes with default parameters (Bankevich et al., 2012). The mitochondrial genomes obtained from shotgun sequencing data were assembled with one of the three following methods: 1) MetaSPAdes with default parameters (Nurk et al., 2017); 2) MetaSPAdes with default parameters followed by the iterative mapping-reassembling approach implemented in TCSF and IMRA (Kinjo et al., 2015) and using the mitochondrial genome of *Cryptotermes brevis* as a reference; C) Novoplasty organelle genome assembler (Dierckxsens et al., 2016) with default parameters and using *C. brevis* mitochondrial genome as a seed sequence. The mitochondrial genomes assembled in multiple contigs were merged using published kalotermitid mitochondrial genomes as references. Assembly gaps were filled with the symbol N (Table S1). All mitochondrial genomes were linearized using MARS (Ayad and Pissis, 2017) to start with the tRNA gene coding for isoleucine and end with the 12S rRNA gene. Each mitochondrial genome linearized with MARS was aligned with MAFFT v.7.475 (Katoh and Standley, 2013) and visually inspected for the presence of 5’- and 3’-terminal assembly artifacts, including: 1) stretches of nucleotides that prevented linearization and 2) duplicated sequences present at both 5’-end and 3’-end of the original contigs. All detected artifacts were removed. We also removed the non-coding portion of mitochondrial genomes between 16S ribosomal RNA and tRNA-Ile which consists mostly of the control region since the sequence repeats in the control region generally leads to artifacts during short read *de novo* assembly. The resulting trimmed mitochondrial genomes were annotated with MitoZ (Meng et al., 2019). The genome annotations of previously published mitochondrial genomes were retrieved from GenBank. MitoZ annotations were visually inspected by first aligning mitochondrial genomes with MAFFT as described above, and then importing the alignments and the MitoZ-generated feature annotations into JalView (Clamp et al., 2004). The errors in MitoZ-generated annotations, mostly comprising of 1) omitted tRNA-Val and 2) 5-terminally truncated 16S rRNA gene were manually corrected. The final mitochondrial genomes and their annotations are available on GenBank under accession numbers <WILL BE UPDATED PRIOR TO PUBLICATION>. Mitochondrial genomes prior to trimming of non-coding region between 12S rRNA and tRNA-Ile are available at <WILL BE UPDATED PRIOR TO PUBLICATION>.

### 18S and 28S annotation

We used BLASTn (Altschul et al., 1990) to search for the 18S and 28S nuclear rRNA genes in the 57 whole-genome shotgun assemblies generated with MetaSPAdes to assemble mitochondrial genomes. The 28S rRNA sequences of *Glossotermes oculatus*, *Zootermopsis angusticollis*, and *Incisitermes schwarzi* were used as BLASTn search queries (GenBank ids: JN647689.1, AY859614.1, GQ337716.1). For each sample, the longest blast hit was identified and oriented to match the 18S and 28S rRNA genes of *Drosophila melanogaster* (GenBank ids: NR_133550.1 and NR_133562.1, respectively). In order to annotate the 18S and 28S rRNA genes of Kalotermitidae, we aligned the 18S and 28S rRNA genes of Kalotermitidae and *D. melanogaster* using MAFFT v.7.475 (Katoh and Standley, 2013). Kalotermitidae 18S and 28S rRNA sequences were then extracted from the multiple sequence alignments. The ribosomal 18S and 28S gene sequences are available on GenBank under accessions <WILL BE UPDATED PRIOR TO PUBLICATION>.

### Alignment and concatenation of the final sequence dataset

The final sequence dataset consisted of 165 mitochondrial genomes and 52 sequences of 18S and 28S rRNAs of Kalotermitidae sequenced in this study. We also used previously published sequences, including the mitochondrial genomes of eight species of Kalotermitidae (Cameron et al., 2012; Bourguignon et al., 2015) (Table S1), the COII sequence of *Ceratokalotermes spoliator* (Thompson et al., 2000), the only described species of the genus *Ceratokalotermes* that was missing from our sampling, and 58 mitochondrial genomes of non-kalotermitid termites that we used as outgroups (Bourguignon et al., 2015).

The rRNA and tRNA sequences were aligned using MAFFT v.7.475 (Katoh and Standley, 2013). For protein-coding genes, we first translated the nucleotide sequences into amino acid sequences with the transeq command of the EMBOSS suite of programs (Rice et al., 2000) using the invertebrate mitochondrial genetic code. The protein sequences were then aligned using MAFFT v.7.475, and back-translated into nucleotide sequence alignments using PAL2NAL v.14 (Suyama et al., 2006). All alignments were concatenated with FASconCAT v.1.04 (Kück and Meusemann, 2010). We generated two alternative alignments, one with the 3rd codon positions of protein-coding genes (WITH3RD) and one without the 3rd codon position of protein-coding genes (WITHOUT3RD).

### Maximum likelihood phylogeny

WITH3RD and WITHOUT3RD alignments were used to infer maximum likelihood phylogenies with IQtree v.1.6.7 (Nguyen et al., 2015). IQtree was run with the options “-m MFP+MERGE - rcluster-max 2000 -rcluster 10 -alrt 1000 -bb 1000 -bnni”. Branch supports were calculated using ultrafast bootstrapping (Minh et al., 2013). The best partitioning scheme was determined using ModelFinder (Kalyaanamoorthy et al., 2017) implemented in IQtree v.1.6.7 from a predefined set of partitions, including one partition for each codon position of the 13 mitochondrial protein-coding genes, one partition for each 22 mitochondrial tRNA genes, one partition for the mitochondrial 12S rRNA gene, one partition for the mitochondrial 16S rRNA gene, and one partition for the combined 18S and 28S rRNA genes. The reconstructed trees were rooted using *Mastotermes darwiniensis*, the extant sister lineage of all other termites (Kambhampati et al., 1996).

### Bayesian time-calibrated phylogeny

We estimated time-calibrated phylogenies of Kalotermitidae using the Bayesian phylogenetic software BEAST2 v.2.6.2 (Bouckaert et al., 2014). We used the uncorrelated lognormal relaxed clock model to account for substitution rate variation among branches (Drummond et al., 2006). The alignment dataset was split into six partitions: one partition for every codon position of the combined 13 mitochondrial protein-coding genes, one partition for the combined nuclear 18S and 28S rRNA genes, one partition for the combined mitochondrial 12S and 16S rRNA genes, and one partition for the combined 22 mitochondrial tRNA genes. Time-calibrated phylogenies were reconstructed using two sequence matrices: 1) the full dataset, including the third codon sites of protein-coding genes (WITH3RD), and 2) a partial dataset, from which the third codon sites of protein-coding genes were removed (WITHOUT3RD). Each partition was given a GTR substitution model with gamma-distributed rate variation across sites (GTR+G). A Yule speciation model was used as tree prior. We used 15 fossils as node calibrations (Table S6). The fossils were selected based on the criteria described by Parham et al. (2002) (for justification concerning the choice of fossil calibrations, see details below). The fossil ages were used as minimum age constraints, which we implemented as exponential priors on node ages. For each fossil, we determined a soft maximum bound using the phylogenetic bracketing approach described by Ho and Phillips (2009). The MCMC chains were run five times independently for both the WITH3RD and WITHOUT3RD alignments. The chains were run for a total of ∼2 x 10^9^ generations and sampled every 20,000 generations. The convergence of the MCMC chains was inspected with Tracer 1.7.1 (Rambaut et al., 2018). The chains for each dataset were combined with Logcombiner v2.6.2 (Bouckaert et al., 2014). We discarded as burn-in the first 10-30% of generations for each MCMC chain. The trees sampled from the MCMC runs were summarized as maximum clade credibility tree with TreeAnnotator v2.6.2 (Bouckaert et al., 2014). The input xml files for BEAST2 analysis are available as Dataset S3.

### Fossil calibrations

We used 15 termite fossils as minimal age internal calibrations (Table S6). In all cases, we used the youngest fossil age estimates reported on the Paleobiology Database (https://paleobiodb.org/). Note that fossil age estimates can be controversial and alternative age estimates exist (see e.g. Kasinski et al. (2020) for hypotheses on Baltic amber age). However, the widths of the 95% HPD intervals produced by our Bayesian time-calibrated phylogeny typically exceed the interval of alternative estimates of fossil ages and we therefore presume that our inference of time-calibrated phylogeny is robust with respect to the use of alternative fossil age estimates. For reproducibility, we adhered here to the ages reported in Paleobiology Database although these might not always reflect the latest consensus on the fossil age. We used the age of *Melqartitermes myrrheus* as the minimal age of all termites (Engel et al., 2007b). *M. myrrheus* is part of the *Meiatermes*-grade that comprised Cretaceous termite fossils intercalating between Mastotermitidae and other termite families (Engel et al., 2009a, 2016). We used *Cosmotermes multus* to calibrate the nodes corresponding to Stolotermitidae + Hodotermitidae + Archotermopsidae. All castes of *Cosmotermes* are known from several aggregations in amber and have similar morphology to modern Stolotermitidae (Zhao et al., 2020). We used *Proelectrotermes swinhoei* to calibrate the Kalotermitidae + Neoisoptera group.

*Proelectrotermes*, along with the fossil genera *Electrotermes* and *Prokalotermes*, were separated by Emerson (1942) into a distinct extinct kalotermitid subfamily Electrotermitinae and considered sister to all other Kalotermitidae, principally owing to what he intuited to be the plesiomorphic retention of tetramerous cerci. As noted by Krishna (1961), these fossils have the typical dimerous cerci of Kalotermitidae and the interpretation of additional cercomeres was an error. *Proelectrotermes* in observable details of the alate imago is diagnosed solely by plesiomorphies (Engel, pers. obs.), and therefore could represent a group outside of crown-Kalotermitidae, being a member of the stem group or is basal to one or more of the earliest-diverging branches. Indeed, in the phylogenetic analyses of Engel et al. (2009, 2016), *Proelectrotermes* was found to be sister to crown-Kalotermitidae, although the available characters were limited.

The phylogenetic placement of *Electrotermes* was intuited by Krishna (1961) to be near *Postelectrotermes* and *Neotermes* based on an assumed transition series in the reduction of mesotibial spines, which were assumed to be symplesiomorphic, and changes in the sclerotization and position of M-vein in the forewing, the latter two of which are quite homoplastic across Kalotermitidae. The mesotibial spines are likely independent in the fossil genera *Electrotermes* and *Proelectrotermes*, as well as modern *Postelectrotermes*, rather than stages of a transition series (Engel, pers. obs.). Indeed, the shortened wing R-vein of *Electrotermes* is more similar to that of African *Glyptotermes* or some *Kalotermes* than to either *Proelectrotermes* or *Postelectrotermes*, particularly the former in which the vein is quite elongated, common across many kalotermitid genera and likely plesiomorphic. Although *Electrotermes* fell outside of crown-Kalotermitidae in recent analyses (Engel et al., 2009a, 2016) the placement is potentially spurious as the genus lacked most codings for the character-state matrix. The genus *Prokalotermes* is a genus of Kalotermitidae known from one species at the Eocene-Oligocene boundary of Florissant, Colorado and another from the Early Oligocene (32 Ma) of Montana (Krishna et al., 2013). While certainly a kalotermitid, the genus remains too poorly understood to provide a more refined phylogenetic placement (Engel, pers. obs.). For these reasons, we did not use *Electrotermes* or *Prokalotermes* as a fossil calibration for crown Kalotermitidae.

The fauna of kalotermitids from the Early Miocene of New Zealand represents a distinctive fauna in ancient Zealandia (Engel and Kaulfuss, 2017). These taxa are poorly understood with unknown affinities to crown kalotermitids. Other Miocene fossil termites, such as those kalotermitids reported from Shandong, China are poorly preserved and documented, and of entirely uncertain taxonomic affinities. Such species are best considered *incertae sedis* and cannot be used as definitive evidence of those genera to which they have historically been assigned. Other fossils of Kalotermitidae include two species in mid-Cretaceous (99 Ma) Kachin amber of the genera *Kachinitermes* and *Kachinitermopsis* (Engel et al., 2007b; Engel and Delclòs, 2010). Both genera, even more so than *Proelectrotermes*, exhibit only plesiomorphies for Kalotermitidae and are likely stem groups. Both are known from only fragmentary remains and are lacking a considerable number of critical traits that would help to better refine their phylogenetic affinities. Thus, both have to be considered as *incertae sedis* among stem-group Kalotermitidae. Trace fossils of kalotermitid nests are known from the Cretaceous through the Miocene, but aside from indicating the presence of kalotermitids with their distinctive frass, nothing more can be determined of the particular groups of Kalotermitidae that produced these ichnological remains.

Note that another fossil species from the Burmese amber, *Archeorhinotermes rossi* (Archeorhinotermitidae), has alates with a fontanelle and is therefore the earliest-known representative of Neoisoptera (Krishna and Grimaldi, 2003), the extant sister group of Kalotermitidae, and could be used to calibrate the same node. We used the age of *Nanotermes isaacae* as a minimal constraint to calibrate the node corresponding to Termitidae + sister group (*Coptotermes* + *Heterotermes* + *Reticulitermes*). *N. isaacae* is the oldest known Termitidae, with unclear affinities to modern subfamilies, but is clearly affiliated to Termitidae (Engel et al., 2011). Eight of the 11 remaining fossils were used to calibrate internal nodes of Neoisoptera, including three nodes of Rhinotermitidae and five nodes of Termitidae. Within the Rhinotermitidae, *Reticulitermes antiquus* is common in the Baltic amber and is clearly assigned to *Reticulitermes* (Engel et al., 2007a), probably representing a stem group *Reticulitermes*, and we therefore used this fossil to calibrate the node corresponding to *Reticulitermes* + sister group. *Coptotermes sucineus* is known from Mexican amber (Emerson, 1971) and was found in the same amber piece with *Heterotermes*, the paraphyletic genus within which *Coptotermes* is nested (Bourguignon et al., 2016b; Buček et al., 2019). We therefore used the fossil of *C. sucineus* to calibrate *Heterotermes* + *Coptotermes*. Finally, *Dolichorhinotermes dominicanus*, a clear member of the *Rhinotermes*-complex (Schlemmermeyer and Cancello, 2000), was used to calibrate *Dolichorhinotermes* + *Schedorhinotermes*. Note that additional species of *Dolichorhinotermes* are known from Mexican amber (Engel and Krishna, 2007b). Within Termitidae, *Macrotermes pristinus*, described by Charpentier (1843) as *Termes pristinus*, and assigned to *Macrotermes* by Snyder, (1949), was used as a calibration for the node *Macrotermes* + *Odontotermes* + *Synacanthotermes*. The four other fossils of Termitidae were described from Dominican amber: *Constrictotermes electroconstrictus*, *Microcerotermes insularis*, *Amitermes lucidus*, and *Anoplotermes* sensu lato. The first three fossil species belong to modern genera (Krishna, 1996; Krishna and Grimaldi, 2009), and were used as minimal age calibrations for the nodes corresponding to their respective genera + sister groups. While the eleven species of the *Anoplotermes*-group described from Dominican amber were assigned to the genus *Anoplotermes* (Krishna and Grimaldi, 2009), they likely belong to different genera within the *Anoplotermes*-group. The genus *Anoplotermes* is presently a polyphyletic assemblage belonging to the South American *Anoplotermes*-group (Bourguignon et al., 2016c). The group is in great need of a comprehensive revision, which implies that fossil species of *Anoplotermes* cannot be precisely assigned to a modern lineage within the *Anoplotermes*-group, and are better referred to as *Anoplotermes* sensu lato. We used these fossils to calibrate all soldierless Apicotermitinae. The last three fossils were used to calibrate the internal nodes of Kalotermitidae. *Huguenotermes septimaniensis* is known from a wing impression found in French sediments dated from the late Eocene. The wing fossil presents plesiomorphies of the clade comprising *Cryptotermes* and *Procryptotermes* (Engel and Nel, 2015) and was used to calibrate *Cryptotermes* + *Procryptotermes* + sister group. *Glyptotermes grimaldii* was used as a calibration for the node including all species of *Glyptotermes* sensu stricto. *G. grimaldii* was found in Dominican amber and closely resembles one extant sympatric species, *Glyptotermes pubescens* (Engel and Krishna, 2007a). For this reason, we considered *G. grimaldii* as a crown-*Glyptotermes* and used it to calibrate the node that contains all species of *Glyptotermes*. The last fossil we used, *Calcaritermes vetus*, was assigned to *Calcaritermes* by Emerson (1969), who stated that this assignment is probable. We considered *C. vetus* as a stem-*Calcaritermes* and used it to calibrate *Calcaritermes* + sister clade.

We used the age of five termite fossils as 97.5% soft maximum bounds. The age of one of these five fossils was used as soft maximum bounds for each 15 nodes with fossil calibrations (Table S6). The maximum soft bounds were assigned using a combination of phylogenetic bracketing and absence of fossil evidence (Ho and Phillips, 2009). For the nodes calibrated with *M. myrrheus*, *C. multus*, and *P. swinhoei*, the three oldest calibrations used in this study, we used as maximum soft bound *Triassoblatta argentina*, the first representative of Mesoblattinidae (Martins-Neto et al., 2005). We used the first fossil of Neoisoptera, *A. rossi* (Krishna and Grimaldi, 2003), as maximum soft bound for the nodes calibrated with the two rhinotermitid fossils *R. antiquus* and *D. dominicanus*, and for the node calibrated with *N. isaacae*, the oldest known fossil of termitids. *P. swinhoei*, one of the oldest undisputed fossils of Kalotermitidae found in Kachin amber, was used as a maximum soft bound for the nodes calibrated with the three crown kalotermitid fossils, *H. septimaniensis*, *C. vetus*, and *G. grimaldii* (Emerson, 1971). The oldest known fossil of *Heterotermes*, *Heterotermes eocenicus* (Engel, 2008), was used as a maximum soft bound for the nodes calibrated with *C. sucineus*. This is justified by the paraphyly of *Heterotermes* to *Coptotermes* (Bourguignon et al., 2016b; Buček et al., 2019). Finally, we used the age of *N. isaacae* as a maximum soft bound for all the nodes calibrated with the crown-termitid fossils, *A. lucidus*, *M. insularis*, *Anoplotermes* sensu lato, *C. electroconstrictus*, and *M. pristinus*.

We excluded several fossils because of their uncertain or revised taxonomic assignments. *Cratokalotermes santanensis* (PaleoDB occurrence numbers 1024939 and 1108956) was included in the Kalotermitidae by Krishna *et al*. (2013), but Grimaldi *et al*. (2008) and the analysis of Engel et al. (2016) demonstrated that it is not a kalotermitid, instead falling within the *Meiatermes* grade of Cretaceous Isoptera.

*Eotermes* was considered not to be a kalotermitid by Krishna (1961). Several Nearctic fossil specimens were assigned to *Cryptotermes* (PaleoDB collection numbers 930175 and 1134865) but lack formal description (Park and Downing, 2001), which precluded their use as fossil calibrations. We did not use *Neotermes grassei* (PaleoDB collection number 995549), from Eocene deposit in France, because its assignment to *Neotermes* was controversial and not accepted by some authors (Nel and Paicheler, 1993; Krishna et al., 2013). In addition, our phylogenetic analyses showed that the genus *Neotermes* is a polyphyletic assemblage, hampering the use of fossils of *Neotermes* as calibrations. Lastly, *Kalotermes piacentinii* (PaleoDB collection number 120351), originally described as a termite fossil (Piton and Théobald, 1937), is no longer considered to be a termite fossil (Nel and Paicheler, 1993).

### Summary evidence time-calibrated phylogeny

A summary evidence time-calibrated phylogeny (SE-tree) was generated by summarizing all phylogenies onto the time-calibrated tree reconstructed with the WITHOUT3RD alignment. Conflicts and supports among trees were summarized using TreeGraph (Stöver and Müller, 2010) and plotted in R. The SE-tree was further summarized to create a reduced SE-tree (RSE-tree) by pruning 36 tips that diverged less than 1 Mya and were collected in the same country.

### Historical biogeography

We reconstructed the historical biogeography of Kalotermitidae using the Bayesian Binary MCMC analysis (Ronquist and Huelsenbeck, 2003) implemented in RASP4.2 (Yu et al., 2015, 2020). The maximum number of areas allowed for each node was set to 1. We ran the model on the two maximum likelihood trees and the two Bayesian time-calibrated trees generated in this study. Each tip was assigned to the biogeographic realm corresponding to its sampling location. We recognized 10 biogeographic realms: Afrotropical, Australian, Madagascan, Nearctic, Oceanian, Oriental, Palearctic, Saharo-Arabian, and Sino-Japanese, as defined by Holt *et al*. (2013), and Neotropical, as defined by Wallace (Wallace, 1876). The analyses were run on each tree with four models: JC (state frequencies set to fixed and among site variation set to equal), JC+GAMMA (state frequencies set to fixed and among site variation set to gamma), F81 (state frequencies set to estimated and among site variation set to equal), and F81+GAMMA (state frequencies set to estimated and among site variation set to gamma). Default values were used for other parameters. Therefore, we ran the Bayesian Binary MCMC analyses four times for every four trees, which makes a total of 16 ancestral range reconstructions of Kalotermitidae.

The sampling was unequal among biogeographic realms, especially for the Neotropical realm that was represented by 45 tips in the RSE-tree. The Neotropical realm was reconstructed as the center of origin for many deep nodes in the phylogeny of Kalotermitidae, possibly reflecting a sampling bias toward Neotropical Kalotermitidae. To rule out an effect of this sampling bias on our ancestral range reconstructions, we performed additional RASP analyses using phylogenies of Kalotermitidae for which Neotropical species were randomly subsampled to 22 and 15 tips. The random subsampling was performed thrice; thence, six RASP analyses were run. RASP was run with the JC model. The node probabilities were plotted onto SE-tree and RSE-tree as pie charts using R. A biogeographic realm map was plotted in R using map data downloaded from https://macroecology.ku.dk/resources/wallace/cmec_regions_realms.zip (Holt et al., 2013). The dispersal age interval plots (Figure 2) were generated in R by traversing the phylogeny in Figure 1 from tips to root. When multiple tips belonged to a single genus that diversified within the realm, only one tip for each such genus was retained. During the traversal, two types of nodes were detected: 1) youngest nodes that with high probability (>90%) do not share the realm with that of the extant representative of each Kalotermitidae genera in each realm, and 2) the oldest nodes that share with high probability (>90%) their realm with all of their descendants. The upper limit of 95% HPD age interval of node type 1 and the lower limit of 95% HPD age interval delimitate the high probability dispersal age interval.

Since the oldest fossils of kalotermitids were discovered in Oriental realm we tested a hypothesis of Oriental distribution of crown Kalotermitidae by constraining the range of the last common ancestor of extant kalotermitids to include Oriental realm. The analysis was performed using R package BioGeoBEARS v1.1.2 (Matzke, 2013) with maximum range size set to 3 and using a dispersal-extinction-cladogenesis (DEC) model (Ree and Smith, 2008).

## Data and script availability

Additional data including all trees generated in this study in Newick tree format, R scripts and input data used to generate figures and tables in this article are available online at https://github.com/AlesBucek/Kalotermitidae_biogeography.

## Acknowledgements

We thank the DNA Sequencing Section and the Scientific Computation and Data Analysis Section of the Okinawa Institute of Science and Technology Graduate University, Okinawa, Japan, for assistance with sequencing and for providing access to the OIST computing cluster, respectively.

## Figures, tables, supplementary data

**Table S1. Summary of sample collection and analytical procedure information.**

**Table S2-S5.** Overview of ancestral realm probabilities and their summary statistics calculated for 10 biogeographic realms, for each of 4 combinations of analyses parameters, and for each of four phylogenetic tree topologies used in RASP analysis (Table S2 and S3: Bayesian inference with and without third codon positions, respectively; IQtree_3rd and IQtree_no3rd: Maximum-Likelihood inference with and without third codon positions, respectively). The table encompass all Kalotermitidae tips — i.e., including tips which are <2 Mya distant and which were dropped in the summary tree in Figure 1. Internal tree nodes with RASP probabilities are in the table defined as the last common ancestor of tree tips with labels “tip1ID” and “tip2ID”. Table columns including mean ancestral realm probabilities and their standard deviations for each realm and each node are denoted as “Means” and “SDs”, respectively, along with the name of each realm.

**Table S6.** Fossil calibration points used in molecular clock analysis.

**Table S7.** Kalotermitid fossils from Paleobiology Database used for Figure S4.

**Table S8.** Constructing and nesting behavior data used for Figure 3.

**Data S1.** Phylogenies in Newick format for all maximum-likelihood and Bayesian inference trees.

---

**Figure S1.**
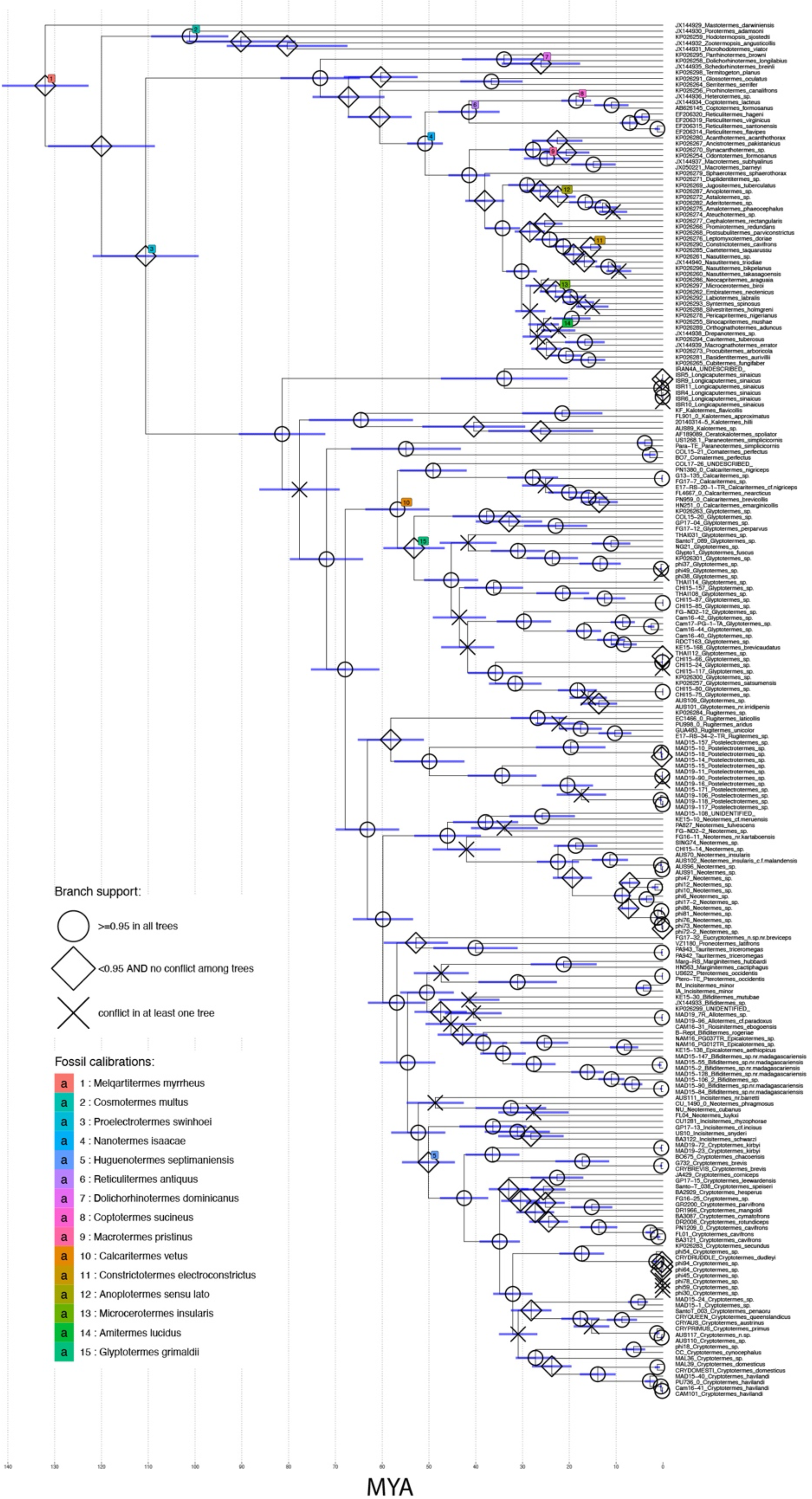
Time-calibrated summary evidence tree (SE-tree) including all Kalotermitidae samples and non-Kalotermitidae outgroups. The tree tip labels consist of sample collection codes (as present also in Table S1) and the species names.

**Figure S2.**
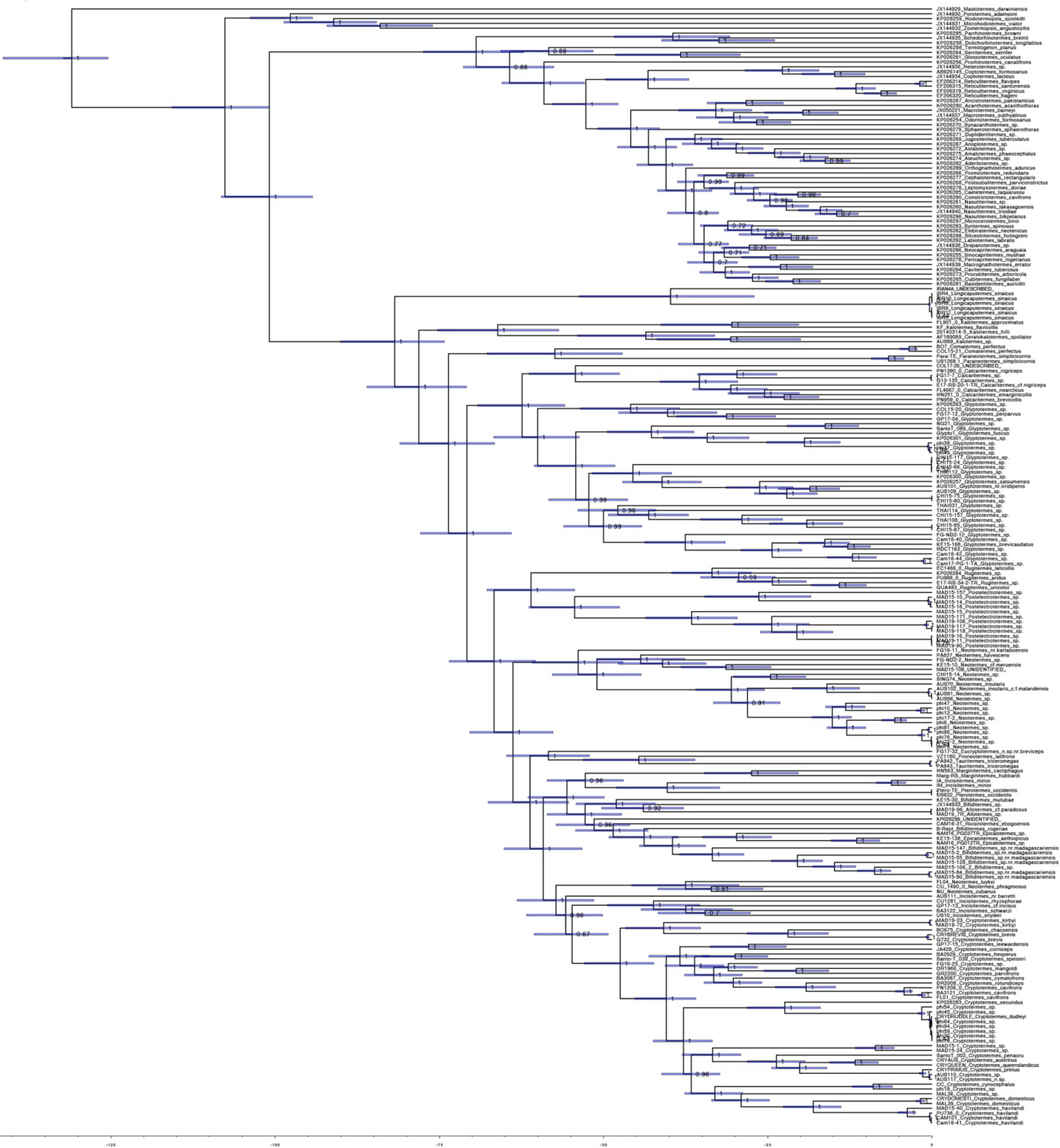
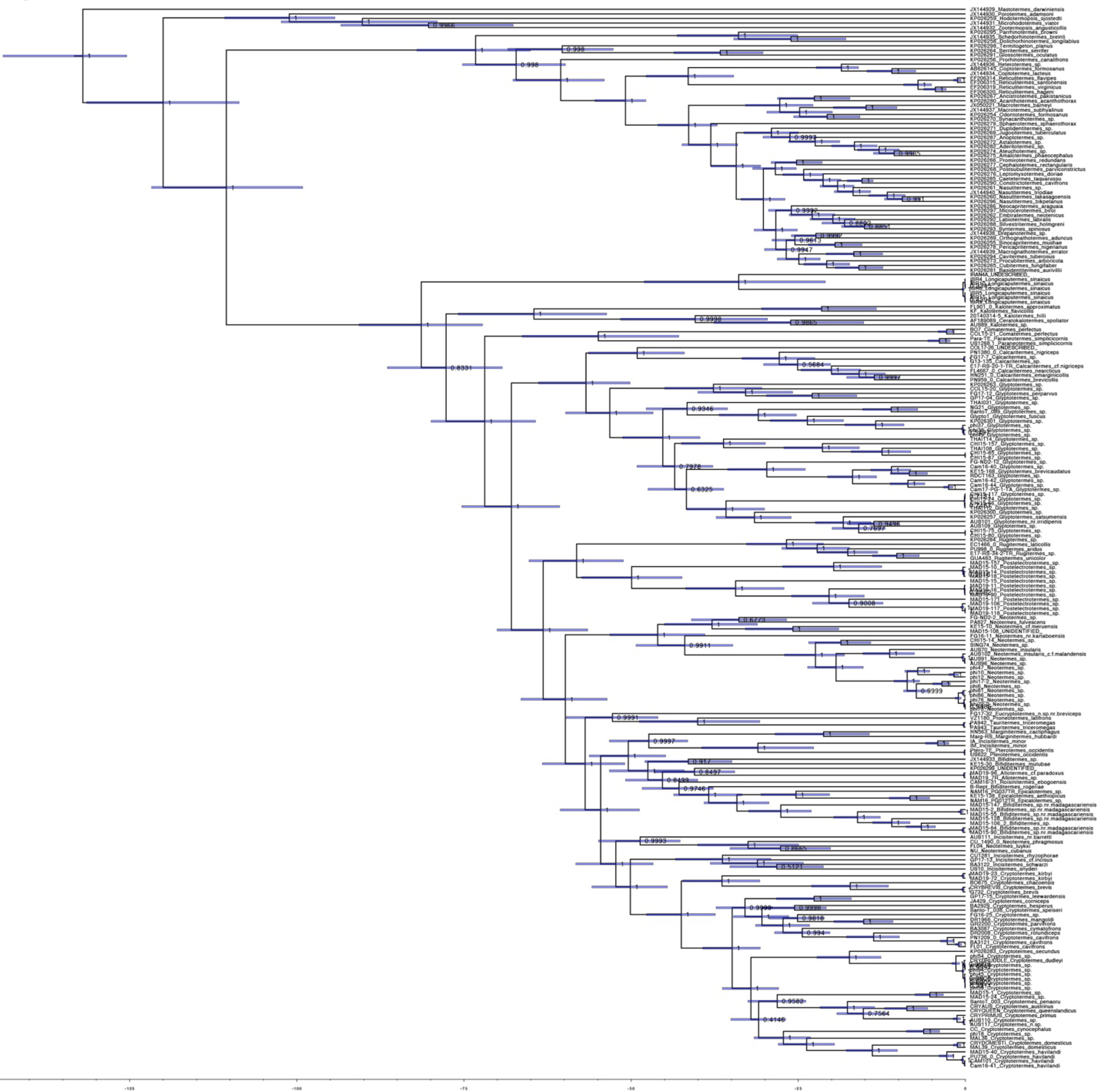
Bayesian phylogenies inferred from sequence alignments (A) with and (B) without third codon positions. Internal node labels indicate posterior probabilities. Node bars delimit 95% HPD intervals. The x-axis time scale is in units of millions of years.

**Figure S3.**
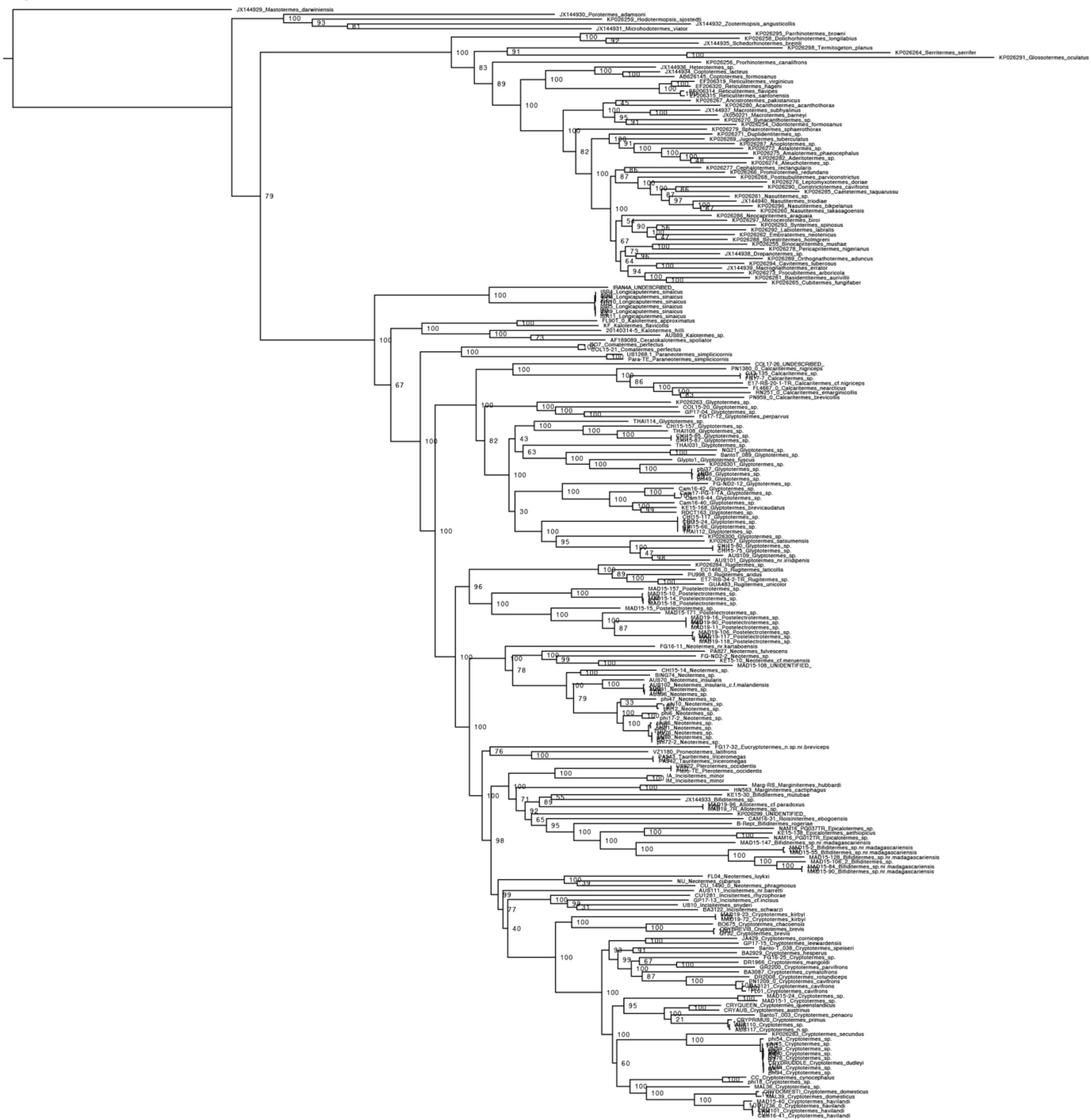
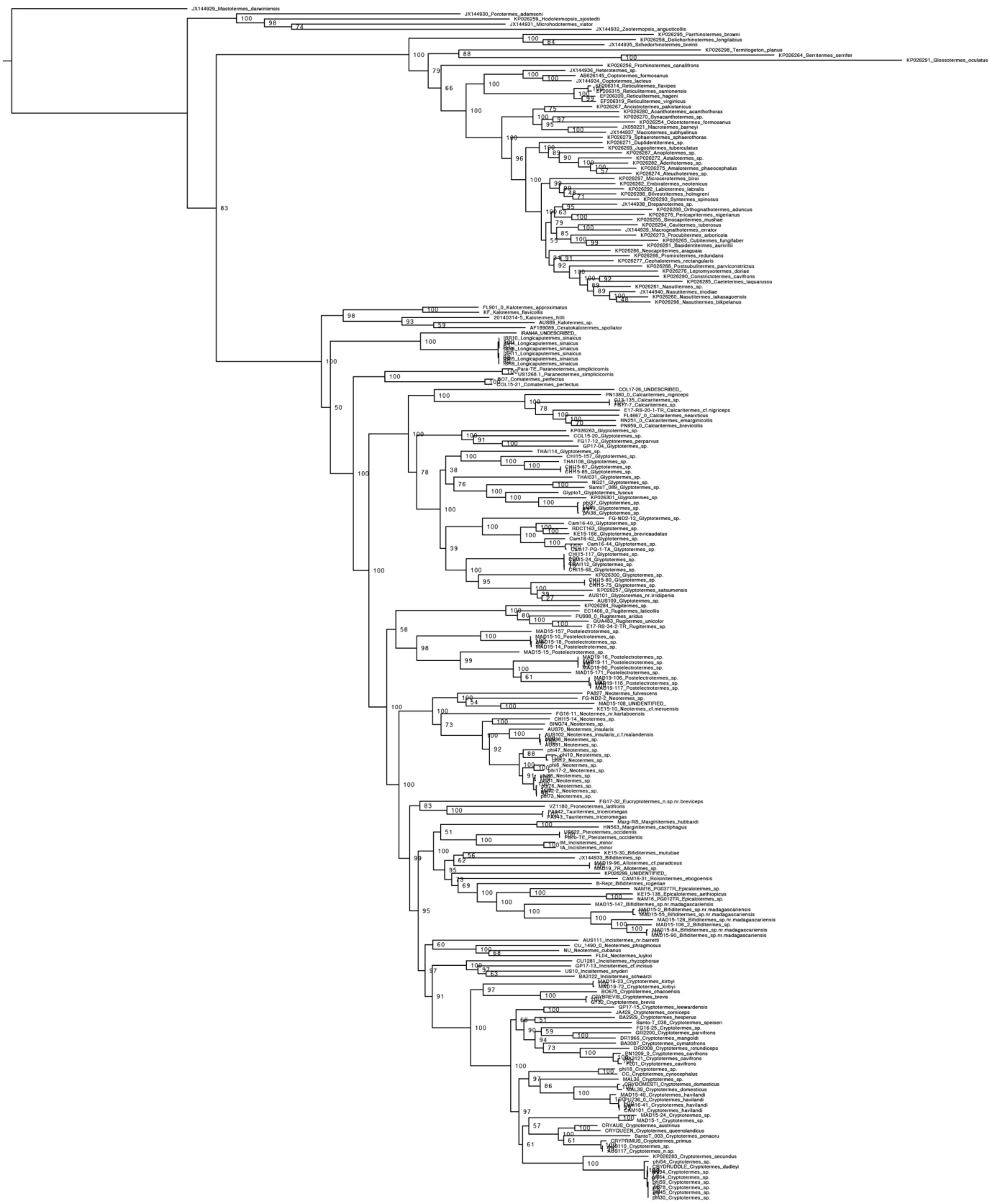
Maximum Likelihood phylogenies inferred from sequence alignments (A) with and (B) without third codon positions. Internal node labels are bootstrap supports estimated from 1000 bootstrap replicates. The trees were rooted by constraining *Mastotermes darwiniensis* as a sister species to the rest of termites.

**Figure S4.**
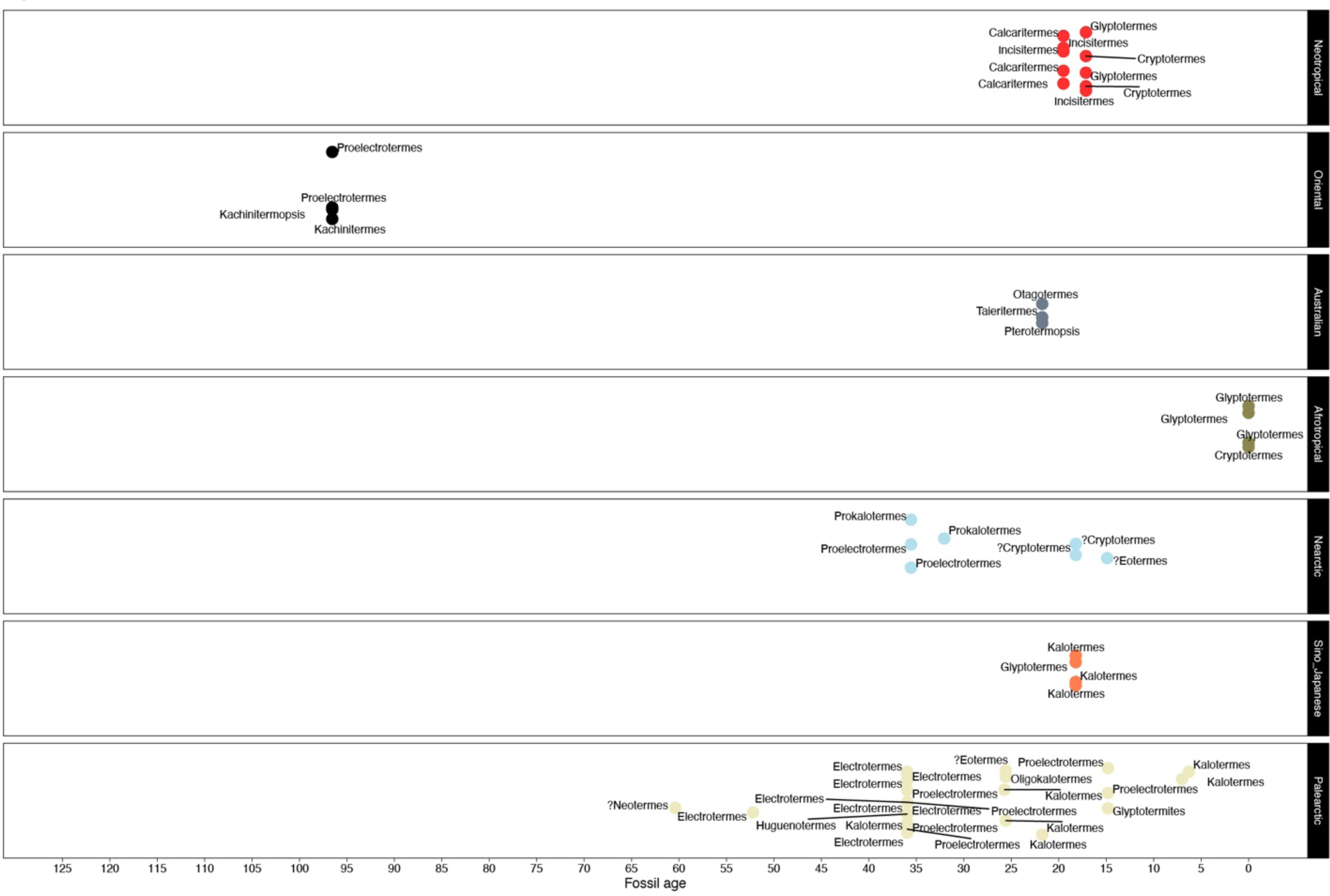
Overview of kalotermitid fossils retrieved from Paleo Database. The fossil records were retrieved from Paleo Database using term “Kalotermitidae” as a database query. The mean ages of fossils are plotted per realm. Interrogative marks prepended to the name of extant genera for which taxonomic assignment of the fossil occurrence is uncertain. Only the taxonomy of fossils that imposed temporal or geographical constraints (i.e., were incongruent) to the historical biogeography reconstructed herein from extant species was revised (see Methods for further details).

**Figure S5.**
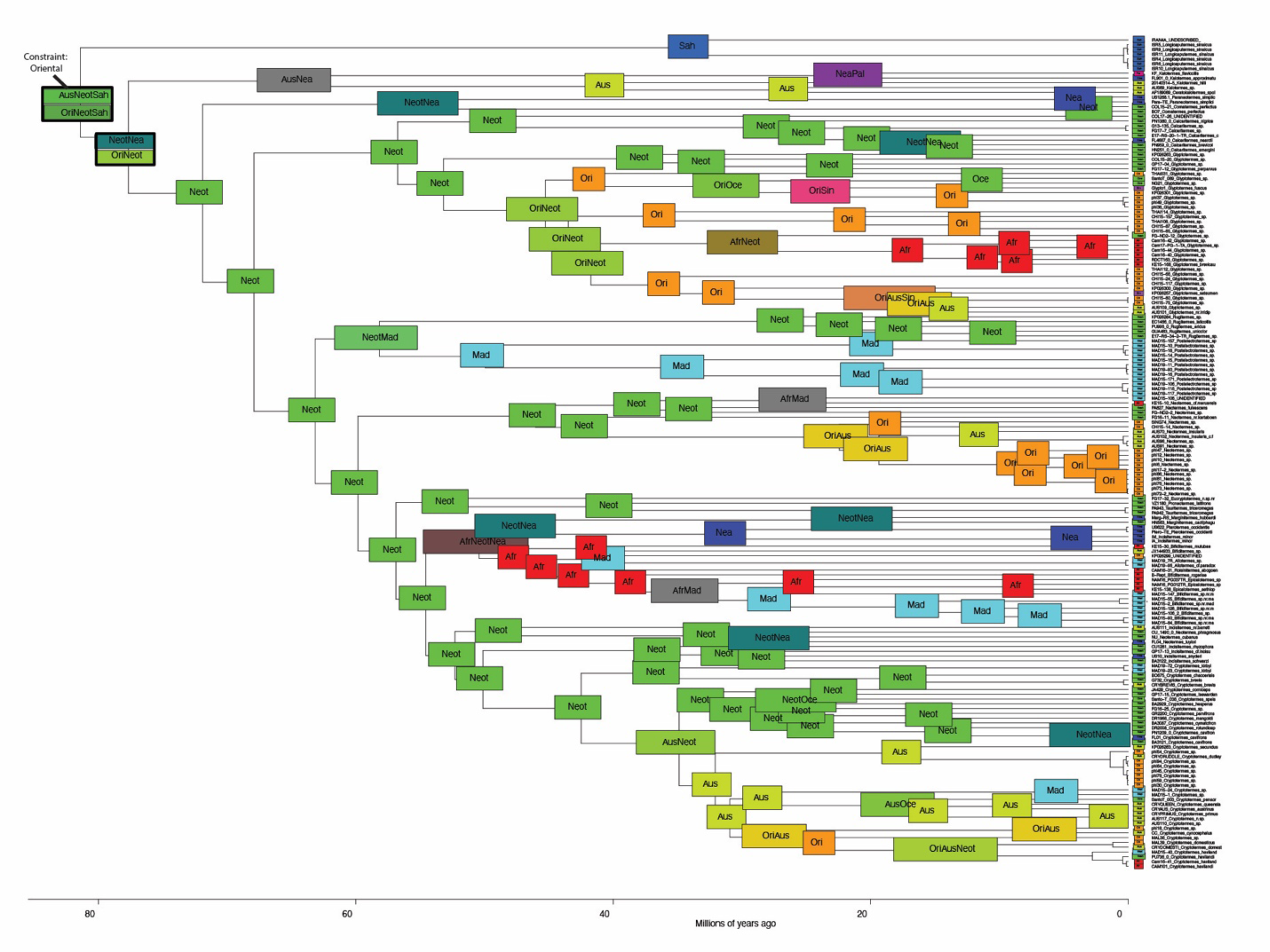
Summary of two historical biogeography inferences under dispersal-extinction-cladogenesis model with and without range of the last common ancestor of extant kalotermitids constrained to include Oriental realm. The ranges with highest probability are shown. Two nodes for which the analysis yielded different range estimates are framed in black and both alternative ranges are shown as two vertically stacked range labels with the upper one showing results of unconstrained analysis and the lower one showing results of constrained analysis. Some states for nodes near the tree tips were omitted for legibility. Realms are indicated by the first three letter abbreviations.

**Figure S6.**
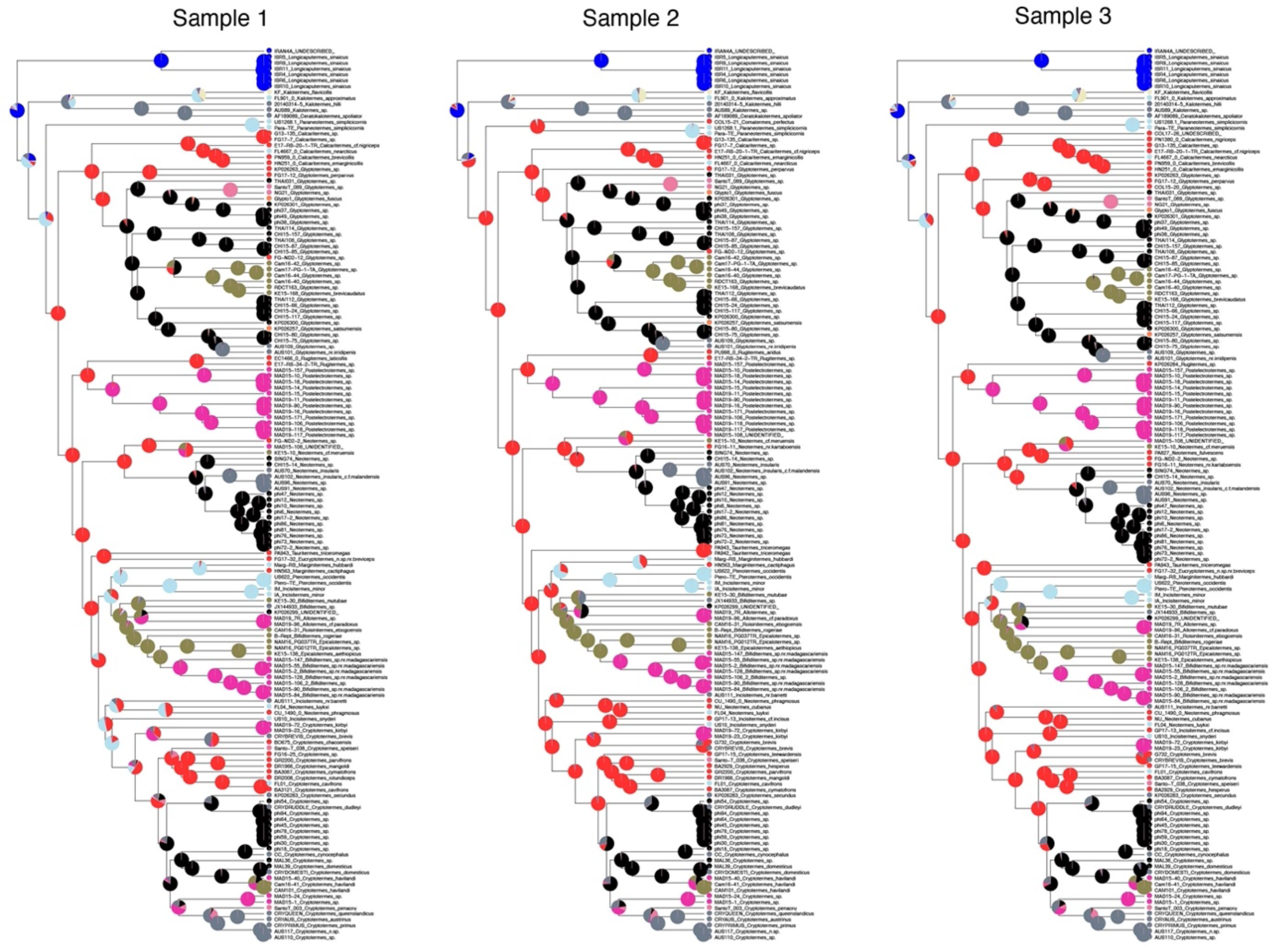
RASP inference of historical biogeography using three sample subsets (Sample 1 - Sample 3) with randomly excluded 23 Neotropical kalotermitid samples.

**Figure S7.**
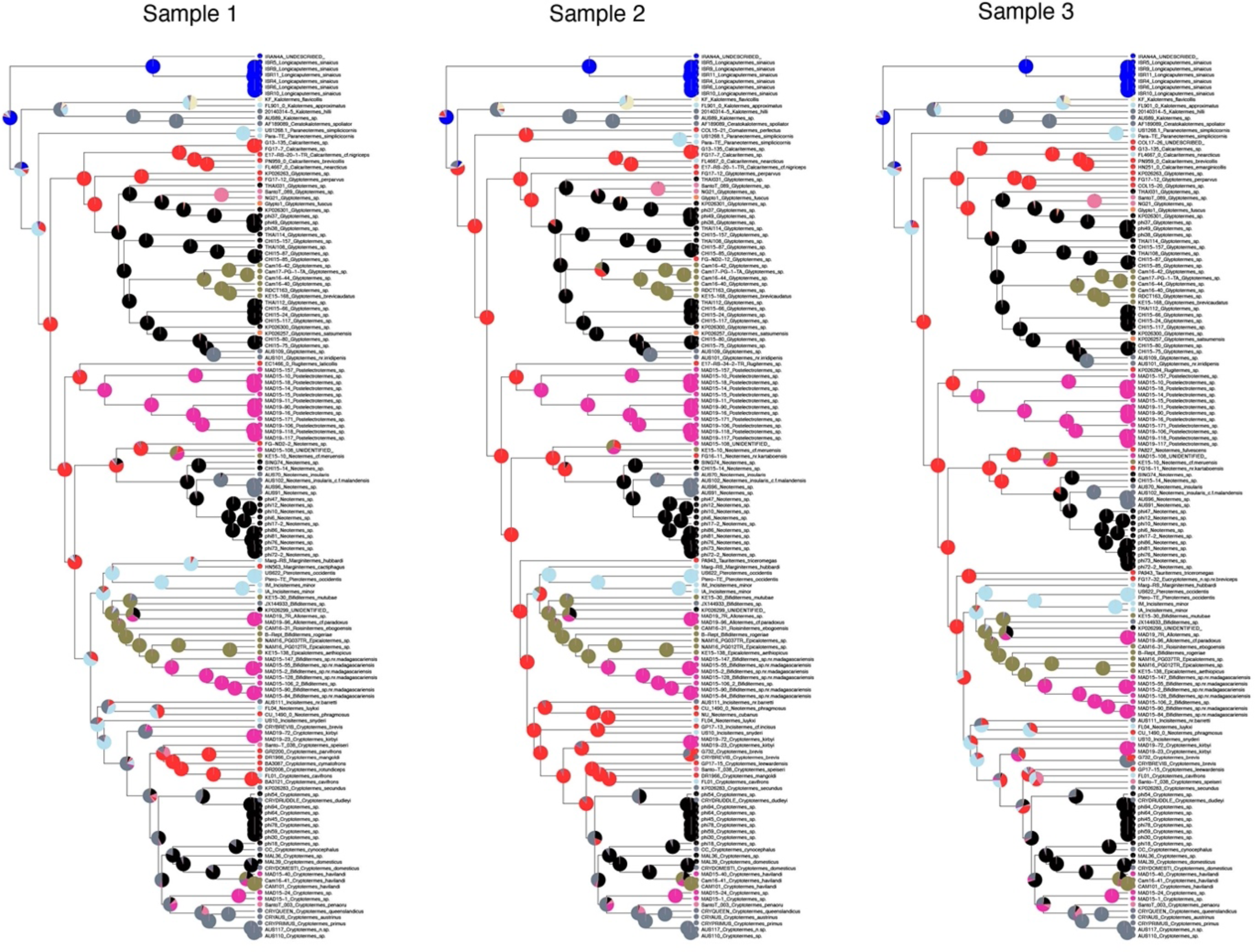
RASP inference of historical biogeography using three sample subsets (Sample 1 - Sample 3) with randomly excluded 30 Neotropical kalotermitid samples.

