## Supplementary figures and images for "Transoceanic voyages of drywood termites (Isoptera: Kalotermitidae) inferred from extant and extinct species"

### Supplemental Figure 1

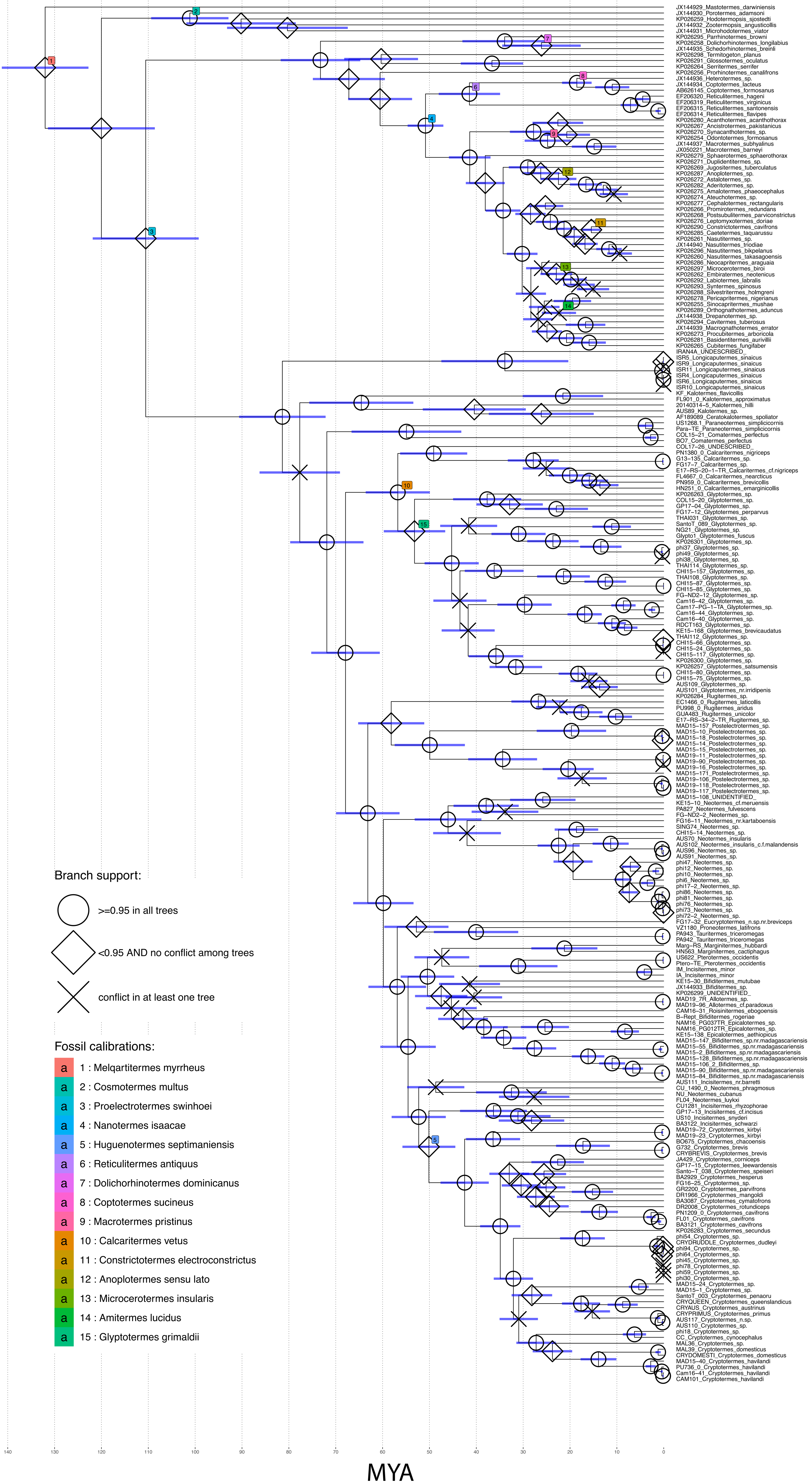

### Supplemental Figure 2A

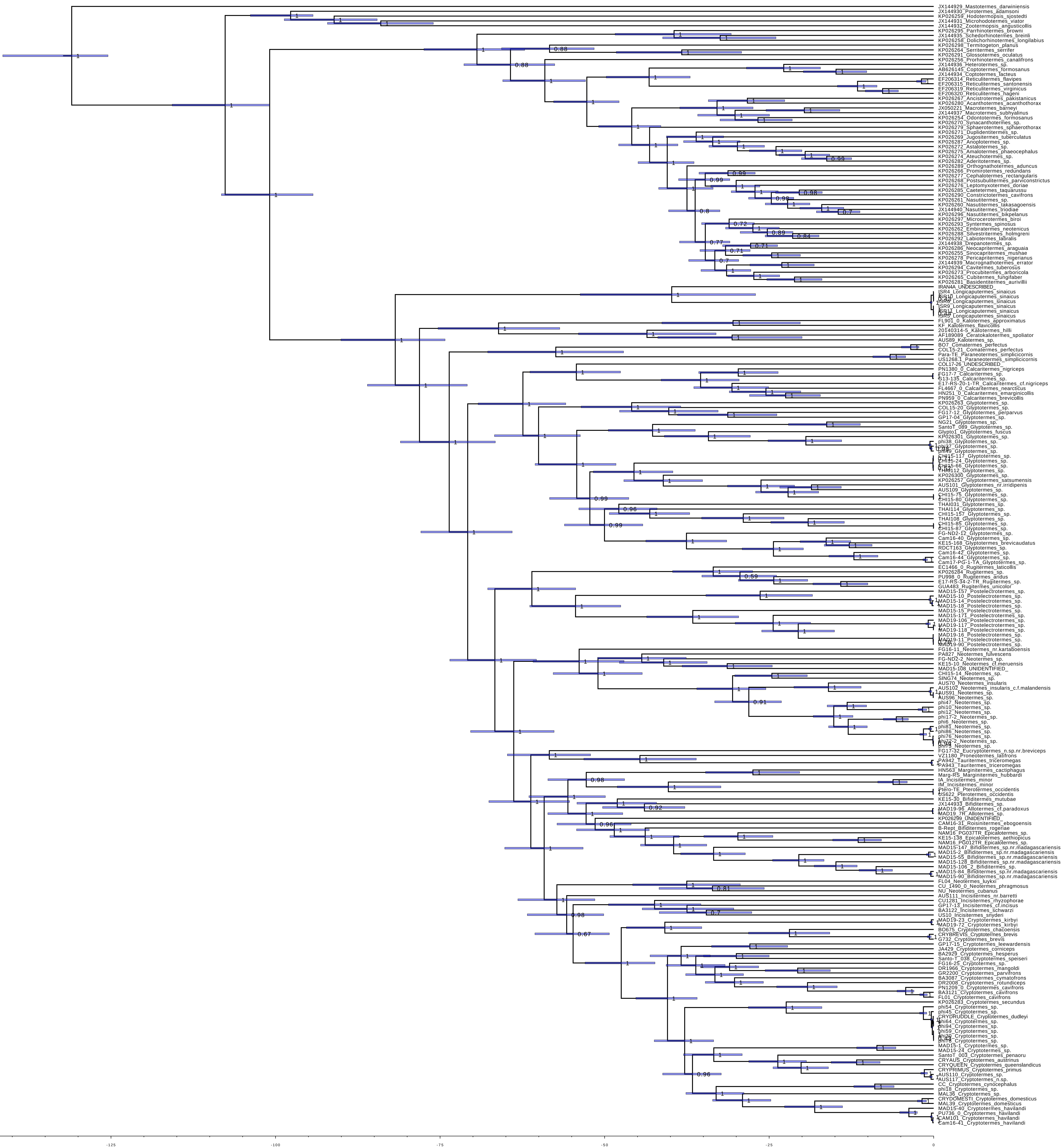

### Supplemental Figure 2B

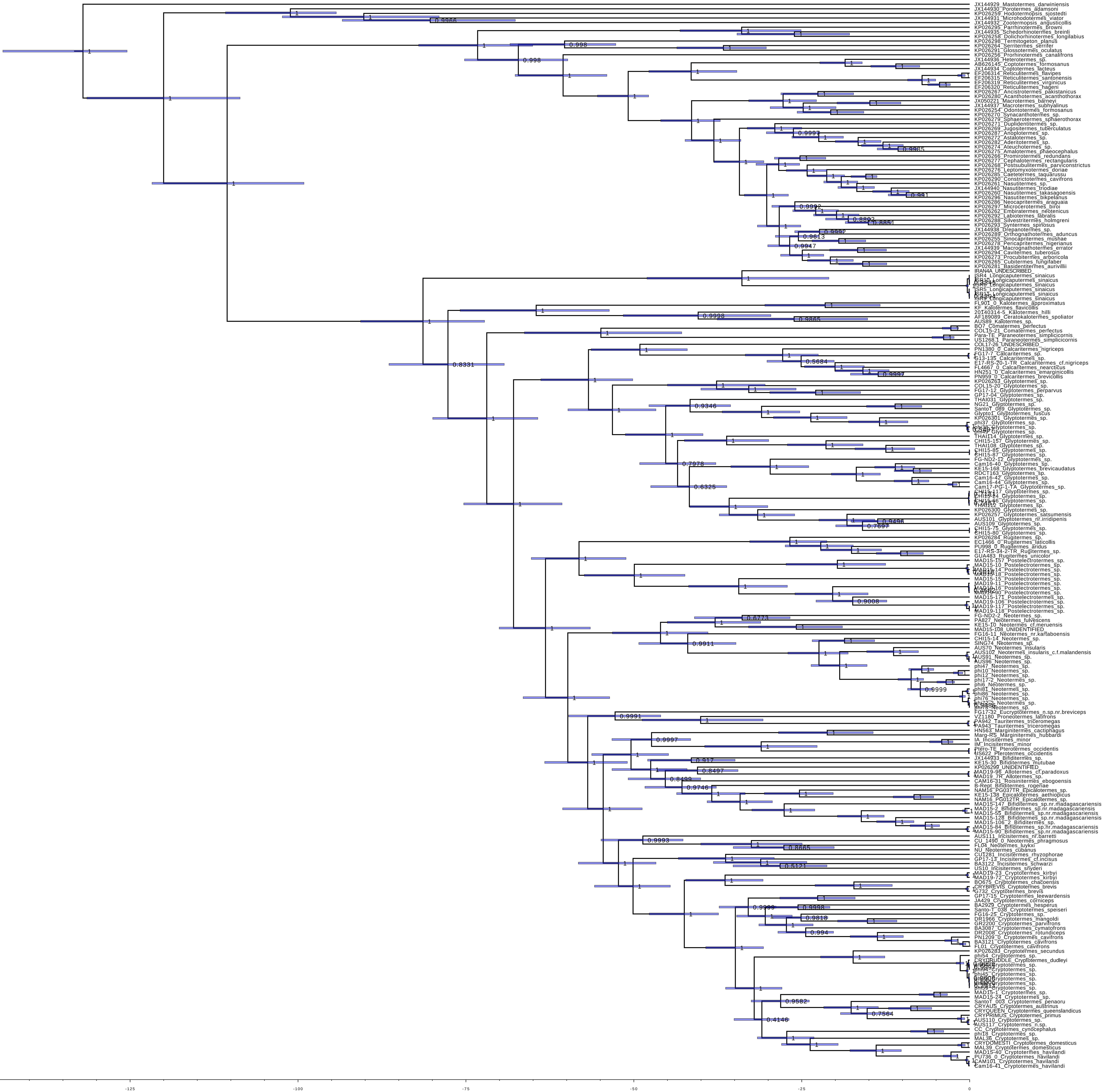

### Supplemental Figure 3A

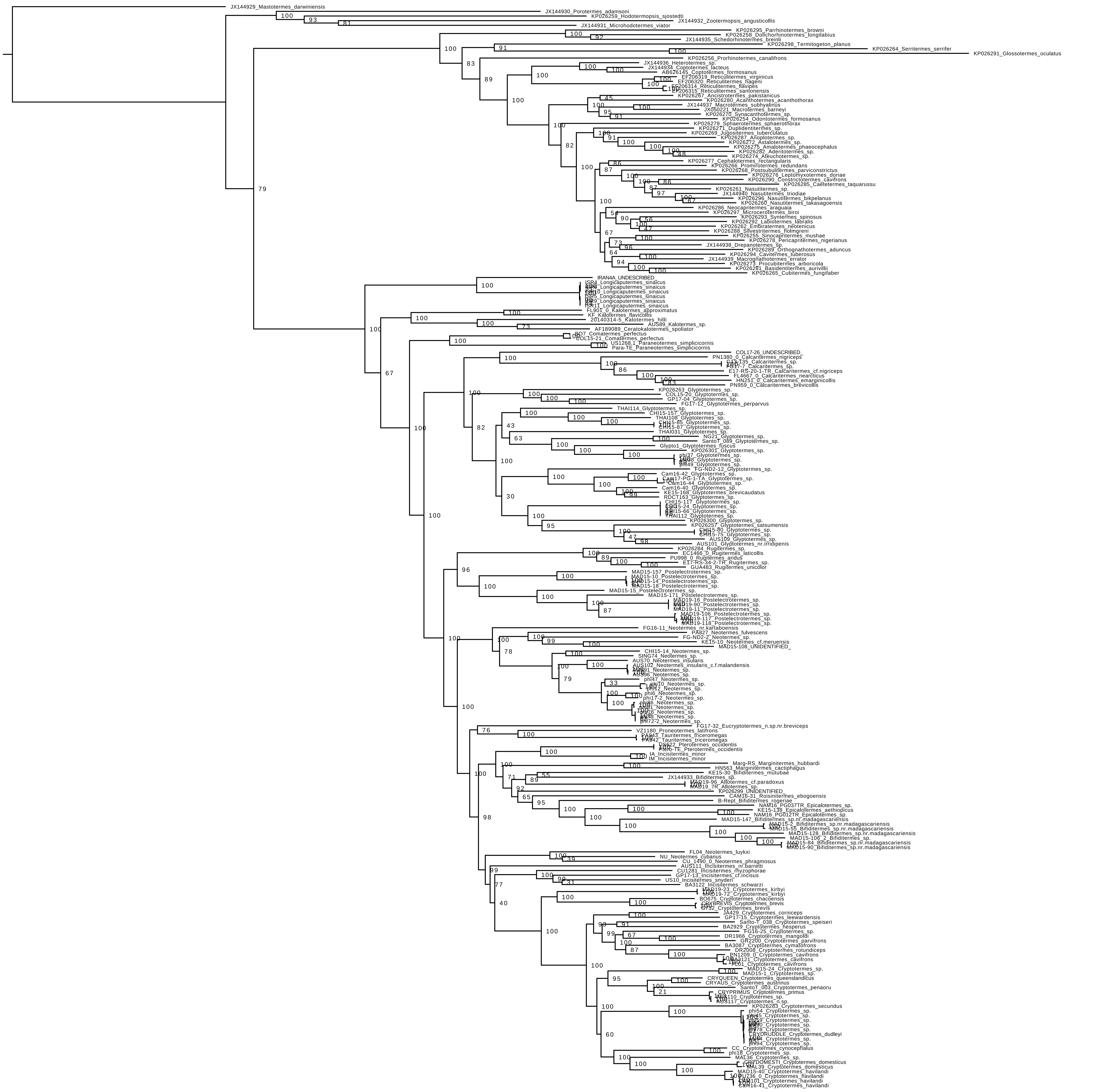

### Supplemental Figure 3B

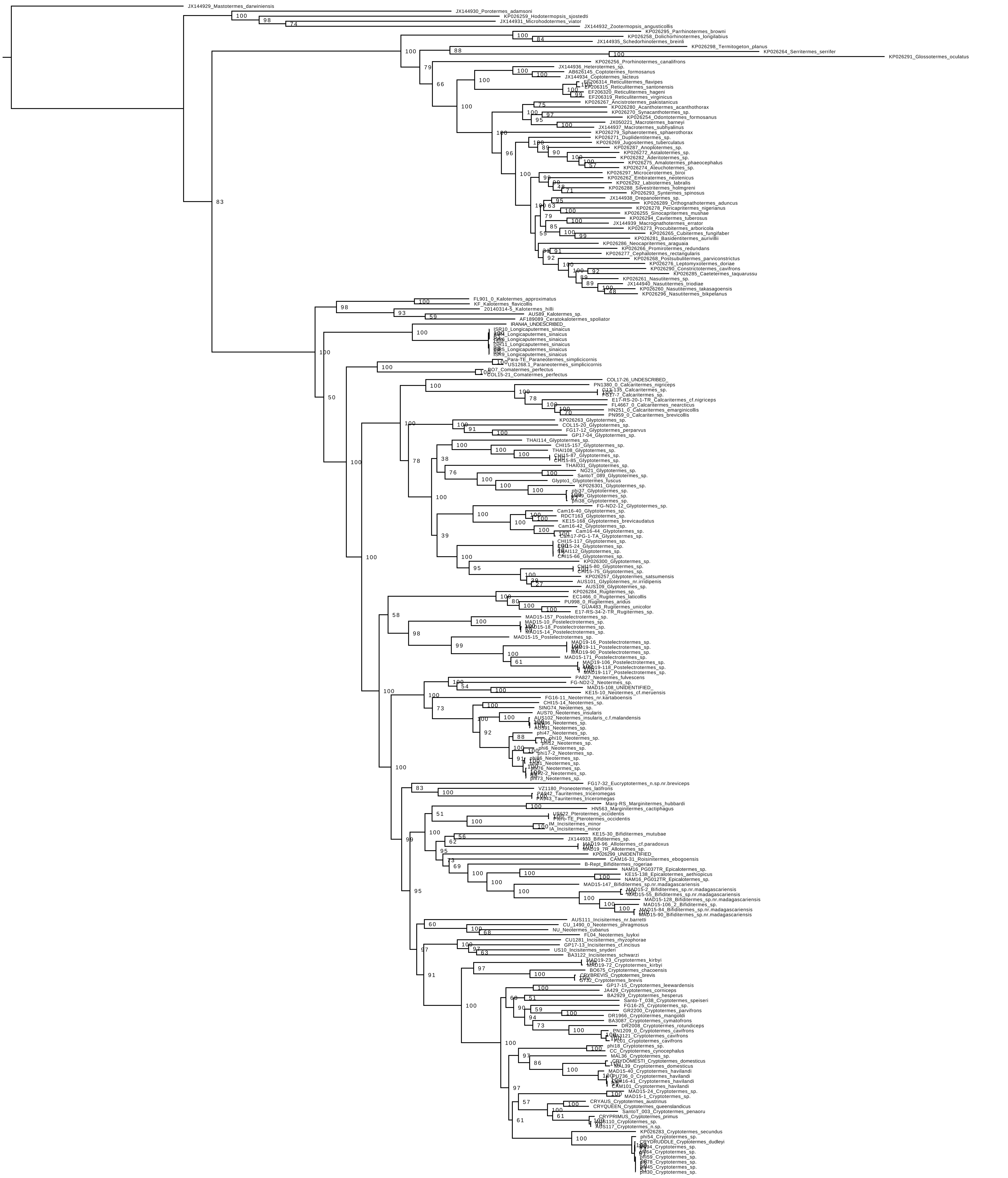

### Supplemental Figure 4

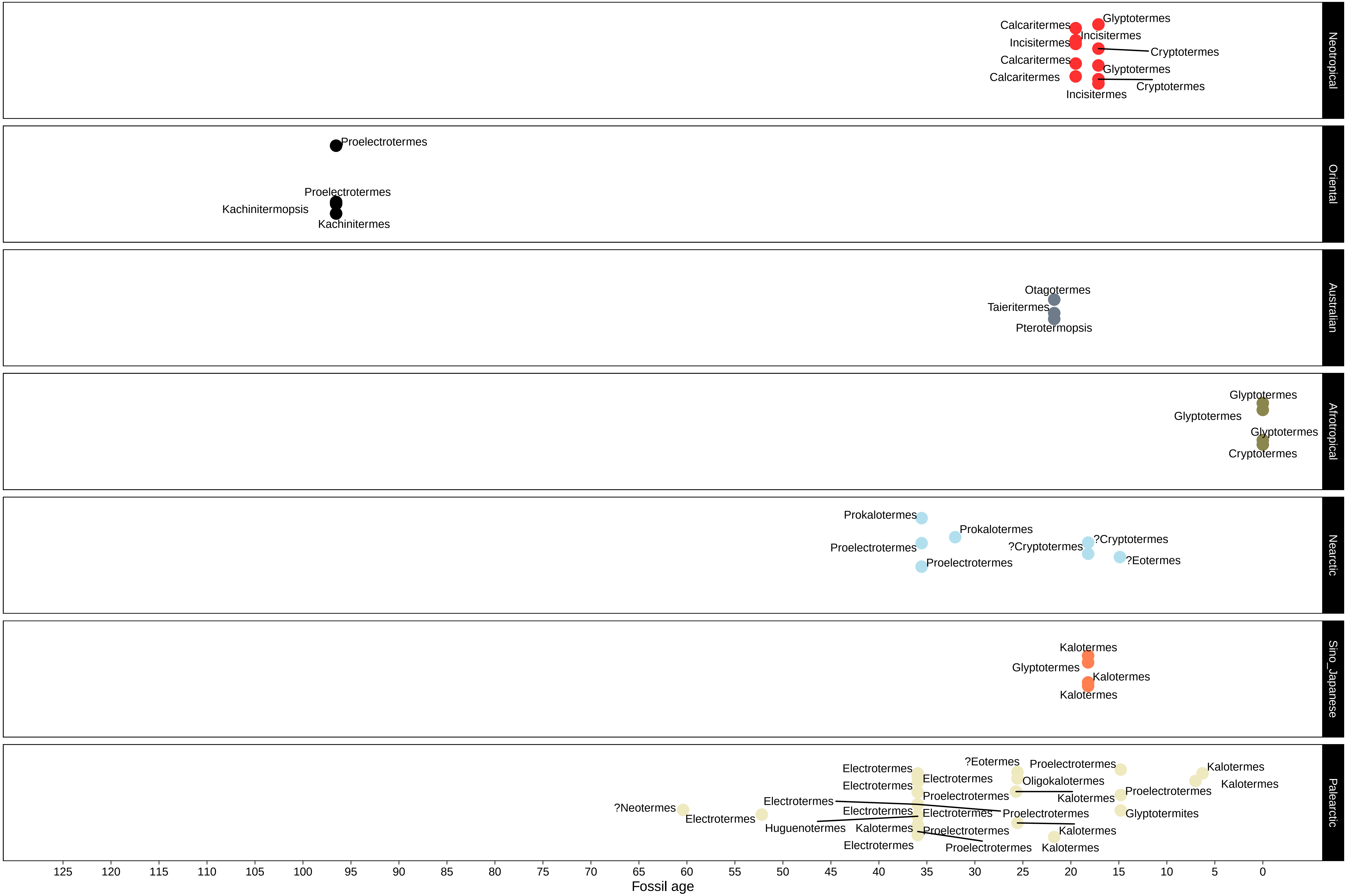

### Supplemental Figure 5

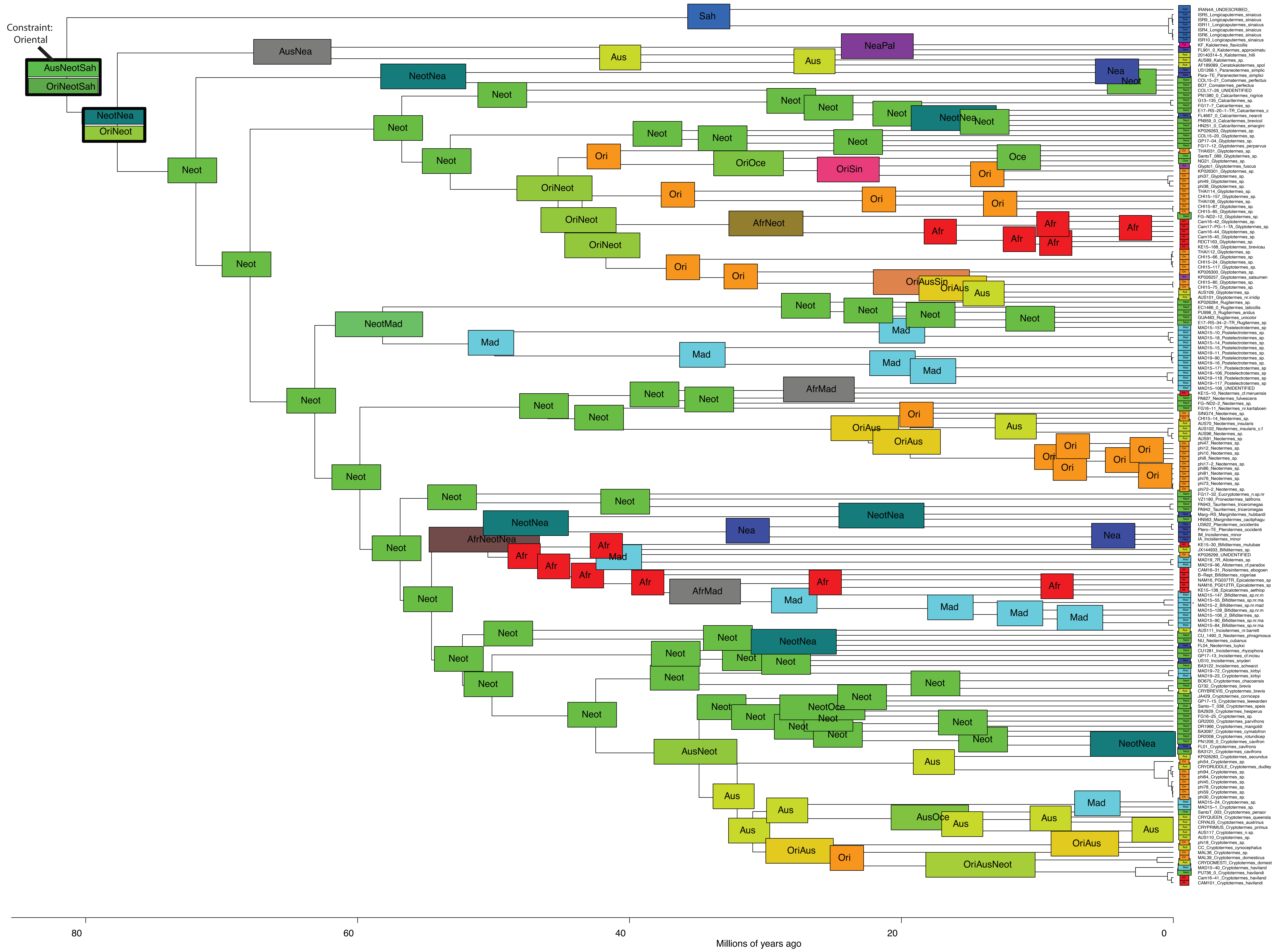

### Supplemental Figure 6

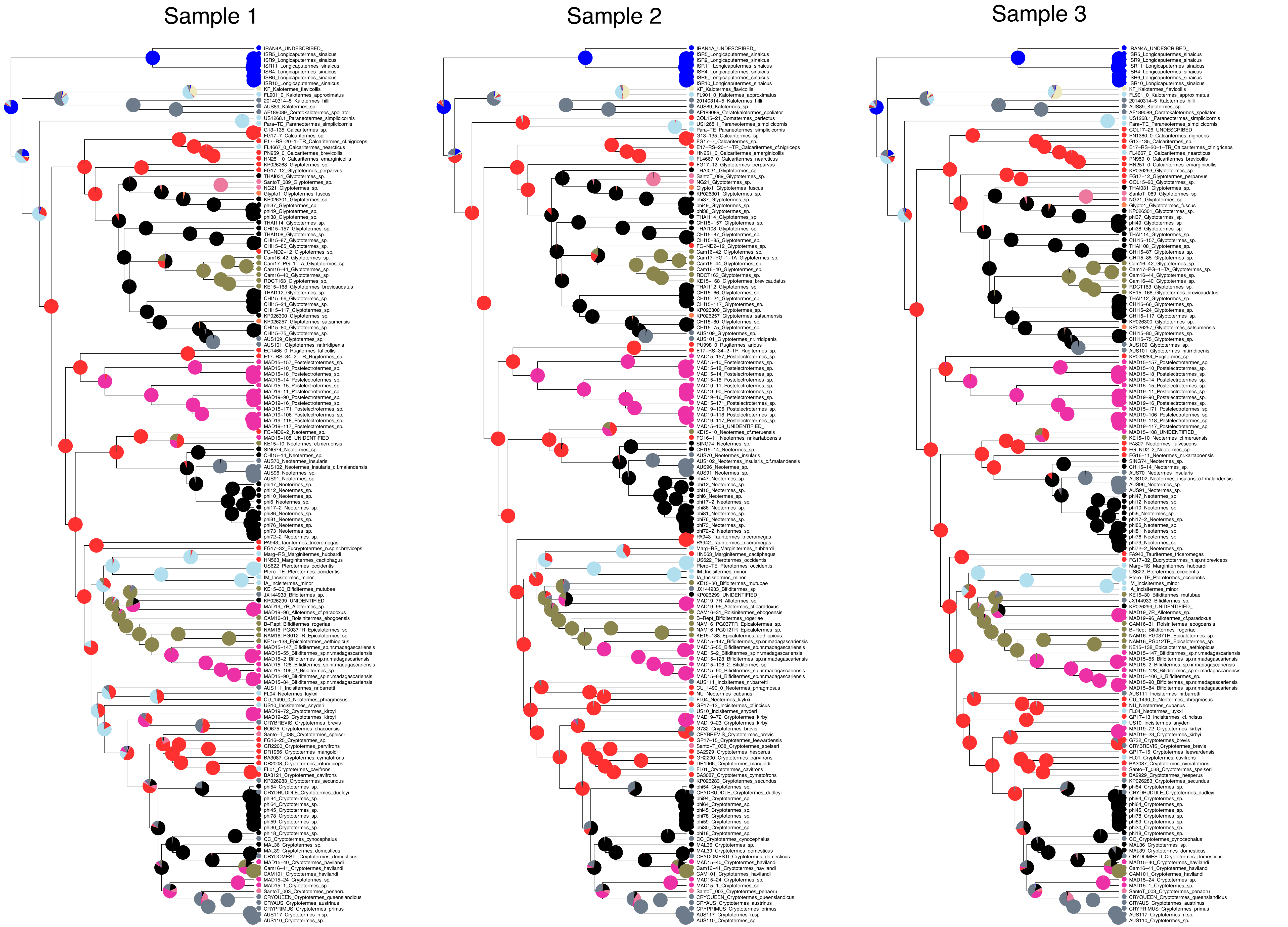

### Supplemental Figure 7

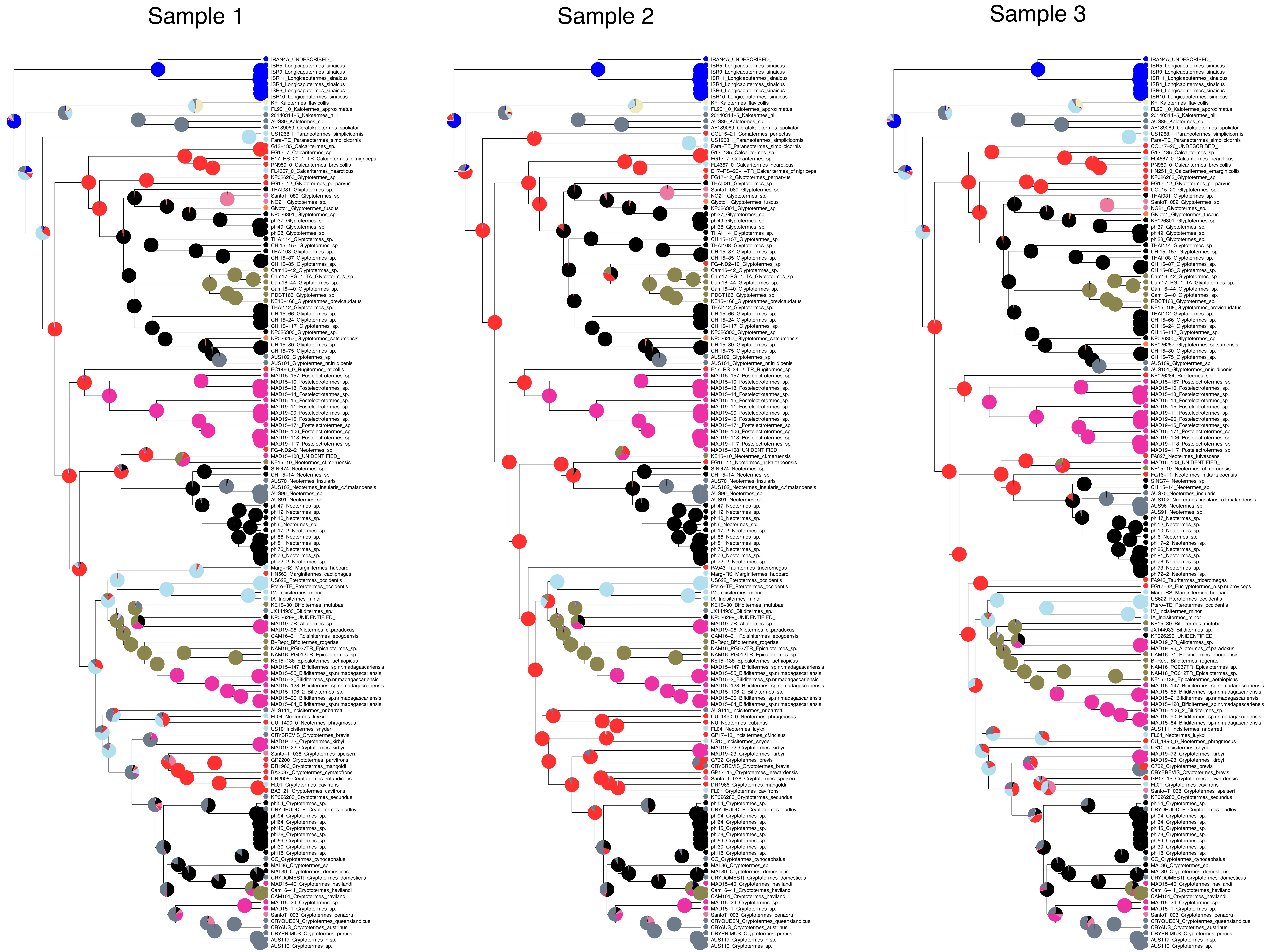
